# Local adaptation of both plant and pathogen: an arms-race compromise in switchgrass rust

**DOI:** 10.1101/2024.06.11.595169

**Authors:** Acer VanWallendael, Chathurika Wijewardana, Jason Bonnette, Lisa Vormwald, Felix B. Fritschi, Arvid Boe, Shelly Chambers, Rob Mitchell, Francis M. Rouquette, Yanqi Wu, Philip A. Fay, Julie D. Jastrow, John Lovell, Thomas Juenger, David B. Lowry

**Affiliations:** Department of Plant Biology, Michigan State University, East Lansing, MI 48824, USA; Great Lakes Bioenergy Research Center, Michigan State University, East Lansing, MI 48824, USA; Department of Horticulture, North Carolina State University, Raleigh, NC, 27607; School of Integrative Plant Science, Cornell University, Ithaca, NY 14850, USA; Department of Integrative Biology, University of Texas at Austin, Austin, TX 78712, USA; Division of Plant Science and Technology, University of Missouri, Columbia, MO 65201, USA; Department of Agronomy, Horticulture and Plant Science, South Dakota State University, Brookings, SD 57006; USDA-NRCS, Kika de la Garza Plant Materials Center, Kingsville, TX 78572, USA; USDA-ARS Wheat, Sorghum, and Forage Research Unit, University of Nebraska, Lincoln, NE 68583, USA; Texas A&M AgriLife Research, Texas A&M AgriLife Research and Extension Center, Overton, TX 75684, USA; Plant and Soil Sciences Department, Oklahoma State University, Stillwater, OK, 74078; USDA-ARS Grassland Soil and Water Research Laboratory, Temple, TX 76502; Environmental Science Division, Argonne National Laboratory, Lemont, IL 60439, USA; Genome Sequencing Center, HudsonAlpha Institute of Biotechnology, Huntsville, AL, USA and the US Department of Energy Joint Genome Institute, Berkeley, CA, USA; Plant Resilience Institute, Michigan State University, East Lansing, MI, 48824

## Abstract

- In widespread species, parasites can locally adapt to host populations, or hosts can locally adapt to resist parasites. Parasites with rapid life cycles locally adapt more quickly, but host diversity, selective pressure, and climatic factors impact coevolution.
- To better understand local adaptation in co-evolved host-parasite systems, we examined switchgrass (*Panicum virgatum*), and its leaf rust pathogen (*Puccinia novopanici*) across a latitudinal range in North America. We grew diverse switchgrass genotypes in ten replicated common gardens spanning 16.78° latitude for three years, measuring rust severity from natural infection. We conducted genome wide association mapping to identify genetic loci associated with rust severity.
- Genetically differentiated rust populations were locally adapted to northern and southern switchgrass, despite host local adaptation in the same regions. Rust resistance was highly polygenic, and distinct loci were associated with rust severity in the north and south. We narrowed a previously identified large-effect QTL for rust severity to a candidate *YSL3*-like gene, and linked numerous other loci to immunity-related genes.
- Both hosts and parasites can be locally adapted when parasites have a lower impact on fitness than other local selection pressures. In switchgrass, our results suggest variation in fungal resistance mechanisms between locally adapted regions.

## Introduction

Local adaptation, the process by which populations within a species adapt to narrow ranges of environmental conditions, is an important force maintaining intraspecific genetic and phenotypic diversity (Kawecki and Ebert 2004). Numerous cases of local adaptation have been described, and researchers have begun to uncover the genetic changes that underlie fitness trade-offs associated with local adaptation (Des Marais, Hernandez, and Juenger 2013; Lowry and Willis 2010; Hall, Lowry, and Willis 2010; Wadgymar et al. 2022; Anderson, Willis, and Mitchell-Olds 2011). However, the majority of local adaptation research has focused on the role of abiotic factors directly on the adapting species, with comparatively fewer studies addressing the role of biotic factors in producing local adaptation (Hargreaves et al. 2020). In host-parasite dynamics in particular, both species may co-evolve local adaptation in an arms-race dynamic, yielding complex evolutionary patterns (Kaltz and Shykoff 1998; Week and Bradburd 2023). Studying the genetic makeup of the co-evolving host and parasite can reveal mechanisms allowing stable coexistence of locally adapted populations within a species.

The balance between host and parasite coevolution is driven by differences in each species’ population and quantitative genetic characteristics. Each may evolve local adaptation to the other, yielding lower infection rates in co-evolved populations if hosts are locally adapted, or higher infection rates in co-evolved populations if parasites are locally adapted (Gandon and Michalakis 2002; Buckling and Rainey 2002; Gandon 2002; Kawecki and Ebert 2004; Greischar and Koskella 2007; Gandon and Michalakis 2002). Parasites are expected to adapt more rapidly, given typically shorter generation times (Gandon and Michalakis 2002). However, migration rates, clonality, and strength of selection play an essential role in determining whether the host or parasite is more likely to be successfully adapted (Greischar and Koskella 2007). Moderate migration in a parasite population can introduce new alleles and speed local adaptation (Gandon 2002), although very high migration rates decrease population barriers and lessen local differentiation (Kawecki and Ebert 2004). Clonal reproduction in parasite populations reduces the effective population size, and may therefore slow the pace of local adaptation (Gandon and Michalakis 2002). Finally, the relative strength of selection plays an essential role. Parasites that prevent host reproduction, such as anther smuts in *Silene*, impose very strong selection for host resistance, so parasite maladaptation can occur (Kaltz et al. 1999). Accurately characterizing host and parasite local adaptation requires a highly-replicated and diverse quantitative genetic experimental design, as well as detailed knowledge of host and parasite ecology.

The traditional standard for diagnosing local adaptation is a reciprocal transplant experiment, preferably with more than two locations (Kawecki and Ebert 2004; Hereford 2009; VanWallendael, Lowry, and Hamilton 2022; Wadgymar et al. 2022). In this experiment, individuals from each environment are transplanted to common gardens both in their local environment and other foreign environments. Metapopulations, sets of populations connected by the movement of individuals, are considered locally adapted when the fitness of populations transplanted to a local environment is higher than foreign populations transplanted to that environment (local-foreign comparison; Kawecki and Ebert 2004). When testing for local adaptation in parasites, researchers typically transplant parasites to different host populations in controlled settings. An alternate method may be used to test for local adaptation in parasites, for which we propose the term “host reciprocal transplant.” Transplanting host genotypes into regions with endemic parasite populations, rather than transplanting parasites between host populations, allows for local adaptation testing that avoids many pitfalls of parasite transplantation (Laine 2007), such as ethical and legal barriers to moving parasites to new regions. Though this method has received less attention in local adaptation theory, it has been used as a strategy in multiple experiments (Davelos et al. 1996; Busby et al. 2014; Cassetta et al. 2023; Laine et al. 2007). However, there are challenges to this method as well. Parasite populations must be both genetically differentiated and consistently present in the transplant locations to detect local adaptation, and the statistical design must account for potential bias from abiotic effects on both the host and parasite (further details in Methods).

Local adaptation studies in parasites often use plant fungal pathogens, owing to their typically narrow host range and economic importance. Fungal pathogens are among the strongest biotic selective forces in wild plant populations, and pose one of the greatest challenges in industrialized agriculture (Fisher et al. 2020). Rust fungi (Pucciniaceae) in particular hold an important place in the history of plant breeding, since introducing genes for stem rust (*Puccinia striiformis*) resistance to commercial wheat was one of the crucial advances of the Green Revolution (Borlaug 1950). The population growth of rusts can depend on both their biotic and abiotic interactions before even reaching the host. Fungi in the Pucciniaceae family often have macrocyclic life cycles, with multiple hosts and five spore-producing forms (Kolmer et al. 2009). For example, the alternate host for wheat stem rust, *Berberis vulgaris*, was brought to North America by colonists before the link to wheat disease was established. It has since been mostly eradicated in the non-native range, aiding stem rust control in wheat (Peterson 2018). The contribution of abiotic conditions to the success of rust infections varies somewhat between species, but generally freezing temperatures, low turbulence, and dry conditions are less conducive to spread and infection (Helfer 2014; Prank et al. 2019). Since all of these conditions are expected to shift with climate change, rust disease may pose greater challenges in the future (Dudney et al. 2021).

Disease resistance is heterogeneously expressed across plant populations, due to historical evolutionary dynamics of hosts and pathogens, as well as interactions with the environment (Thrall and Burdon 2002; Kniskern and Rausher 2006; Chappell and Rausher 2016; Colhoun 2003; Atkinson and Urwin 2012; Huot et al. 2017; VanWallendael et al. 2020). The high fitness cost of fungal disease means that if resistance loci are present, they can sweep rapidly to fixation in plant populations (Bergelson, Dwyer, and Emerson 2001). However, pathogens themselves face a high cost imposed by plant resistance, so mutations that increase infectivity also spread rapidly (McDonald and Linde 2002), contributing to arms-race dynamics. Fitness costs can also come from the mechanism of either resistance in the host or infectivity in the pathogen (Simms and Triplett 1994; Laine and Tellier 2008), though the strength of these costs has been challenging to predict since they may only manifest under certain environmental conditions. Thus, disease response evolution in natural systems is a patchwork of resistance and susceptibility, often dependent on abiotic environmental conditions.

Since the Green Revolution, wheat stem and leaf rusts have been widely researched, offering a view of patterns in resistance loci (McIntosh, Wellings, and Park 1995; Feuillet et al. 1995; Lillemo et al. 2008; Yu et al. 2014). Host plant resistance to fungal pathogens can take multiple forms ranging from resistance from a few immunological resistance loci (Salcedo et al. 2017; Asnaghi et al. 2001) or polygenic resistance that includes structural or life-history traits (Yu et al. 2019). While some wheat stem rust resistance mechanisms such as *LR13* and *LR34* have been effective under diverse growing conditions and against many strains, emergence of novel rust strains has threatened breeding gains (Singh et al. 2011).

Researchers have found that resistance loci vary greatly in their mechanisms of plant protection. Resistance often varies between host life stages, or may be specific to certain rust strains (Kolmer and Liu 2000). While wheat varieties vary in their constitutive resistance to rust, breeders have focused on uncovering single-gene resistance (R-gene resistance) that can be introgressed into susceptible varieties. Mechanistically studied genes include *Sr35*, a wheat gene from Einkorn (*Triticum monococcum*) that confers resistance to the Ug99 stem rust strain (Salcedo et al. 2017) by binding pathogen effectors to trigger activation of the *ZAR1* resistosome (Förderer et al. 2022). Despite rapid advances in understanding mechanisms of rust resistance, knowledge of the degree to which these conclusions are transferable to other rust-infected plants is lacking.

Switchgrass (*Panicum virgatum* L.) is a locally adapted perennial plant conducive to pathogen local adaptation study, since it is consistently infected with several fungal pathogens including rust (*Puccinia novopanici* Demers), leaf pathogens in the *Bipolaris* and *Colletotricum* genera, and the panicle smut *Tilletia maclaganii* (Kenaley et al. 2019). Switchgrass genetic diversity is divided into three major genetic populations that mostly correspond with locally adapted ecotypes. The Midwest population is common in the north-central region, the Atlantic population along the east coast, and the Gulf population in Texas and along the Gulf of Mexico (Lovell et al. 2021). These populations differ greatly in phenotypic traits and in abiotic stress response, potentially owing to adaptive fitness trade-offs. Fitness trade-offs that underlie local adaptation can be caused by genomic effects such as linkage or pleiotropy (Wadgymar et al. 2017). For instance, if a gene that enhances freezing tolerance is linked to a gene that results in disease susceptibility, the genotype would perform well in the cold, but suffer in regions where pathogen pressure is higher, as is often the case for genotypes in the Midwest population. The Gulf and Atlantic populations typically have higher overall resistance to leaf fungi, but this can vary between years (VanWallendael et al. 2020). Previous research in a different common garden system revealed that rust resistance patterns differ greatly between the northern and southern US, though there was no difference in rust species composition across space (VanWallendael et al. 2020). We found large-effect Quantitative Trait Loci (QTL) for rust resistance on chromosomes 3N and 9N, but the outbred mapping population did not have sufficient resolution to narrow these to specific genes.

In this study, we sought to understand regional differences in resistance, and to more closely pinpoint genetic loci associated with rust infection severity using a genome-wide association study (GWAS). Given our understanding of rust abiotic habitat preferences as well as host ecotypes, we expected to find pathogen population differentiation between northern and southern regions. Further, we expected that the shorter generation time of rust than switchgrass will result in it being locally adapted to its switchgrass host, rather than switchgrass to rust. We aimed to recapitulate previous results showing regional differences in rust resistance loci, and identify candidate genes involved in rust resistance. We predicted that resistance would involve large-effect immune genes, similar to those found in wheat. Since resistance is so clearly differentiated across locally adapted switchgrass ecotypes, we expected that it is genetically linked to other traits that differentiate populations, such as biomass production or climatic niche.

## Materials and Methods

### Experimental Design

This study used an experimental design described in Lovell et al. 2021. Briefly, 1347 switchgrass rhizomes were collected from numerous locations in the United States (Figure 1). These rhizomes were grown, then clonally propagated at a central facility in Austin, Texas, then transplanted to field conditions in 2018. Three sites: Austin, TX (PKLE), Columbia, MO (CLMB), and Kellogg Biological Station, MI (KBSM) were planted with all surviving 1218 genotypes, and an additional six sites were planted with a core subset of 630 tetraploid genotypes. The remaining octoploid and hexaploid genotypes were measured for rust and other traits, but not used in the mapping population (Napier et al. 2022). At each site, switchgrass rhizomes were planted in a honeycomb grid with 1.6m spacing between plants. They were watered to promote establishment, and periodically weeded each season, but otherwise received no treatments. We planted a border of short-statured lowland switchgrass (Blackwell cultivar) around the exterior of the plot, and dead plants were replaced with Blackwell “placeholders” to reduce edge effects. The genetic makeup of the core group was assessed using whole-genome resequencing, which yielded over 11 million polymorphic single nucleotide polymorphisms (SNPs) with minor allele frequency > 0.05.

**Figure 1:**
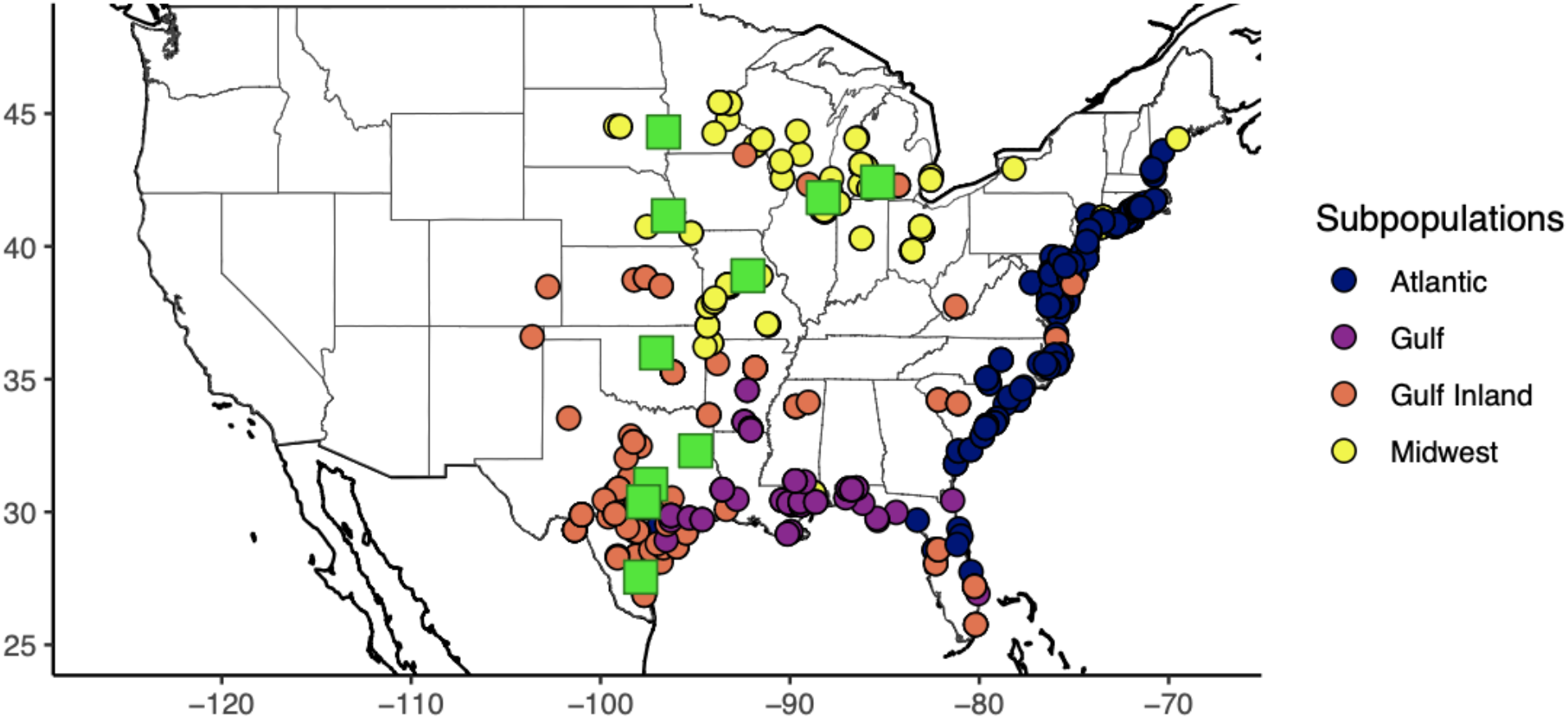
Collection locations (circles), and planting sites (green squares) for the replicated diversity panel. Coloring for circles indicates population membership based on shared SNPs. From north to south, the sites are BRKG (Brookings, SD), KBSM (Kellogg Biological Station, MI), FRMI (Fermilab, IL), LINC (Lincoln, NE), CLMB (Columbia, MO), STIL (Stillwater, OK), OVTN (Overton, TX), TMPL (Temple, TX), PKLE (J.J. Pickle Research Campus, TX), and KING (Kingsville, TX). We were not able to obtain rust ratings from STIL, nor rust population samples from OVTN.

### Rust Population Genetics

In 2019, we collected rust-infected leaves from nine of our site locations to assess pathogen population structure. For most sites, these samples were taken from adjacent(< 500m) switchgrass experimental plots used for a previous mapping study (VanWallendael et al. 2020) which were further-developed at the time, and therefore yielded more rust-infected leaf tissue. At the Fermilab site (FRMI), we sampled from the GWAS diversity panel, since we did not have a previous experimental plot. We haphazardly collected 20 samples showing clear sori from each site. The samples were dried on silica gel and stored at room temperature before processing at Cornell University. We scraped spores from sori on each leaf, then extracted genomic DNA from each sample using a DNeasy Plant Mini Kit (Qiagen, Valencia, CA), following the manufacturer’s protocol with some modifications as described in Kenaley et al. (2016). The samples were then flash frozen in liquid nitrogen and ground separately to a fine powder using a mini-pestle. Next, 500 µL AP1 lysis buffer and 10 µL proteinase K (10 mg mL-1; Qiagen) were added to each tube, and the samples were incubated overnight (55 °C at 1100 rpm) in a thermomixer (model 5350; Eppendorf, Hauppauge, NY). After incubation, the tubes of lysed tissue were placed in boiling water for 5 min, then centrifuged briefly (3000 rpm for 5s), and 150 µL of the neutralization buffer (P3 buffer) was added immediately to each tube. The tubes were then centrifuged (13000 rpm, 1 min) and placed at -20 °C for 10m to pelletize cell debris and ease transfer of the supernatant (∼ 660 mL) to a QIAshredder Mini spin column, tubes were again centrifuged for 5 min at 13,000 rpm. The remaining procedure was identical to that provided in the DNeasyPlant Mini protocol (steps 9-19, Qiagen). After DNA quality and quantity filtering, 88 samples remained with at least 5 samples from each site. We shipped samples to the Texas A&M Genomics and Bioinformatics Service for library preparation and sequencing on an Illumina NovaSeq 6000 with an S4 flowcell. We sequenced the 99.9 Mb genome to ∼50x coverage per sample.

Since *P. novopanici* does not grow in culture, we could not sequence single-spore isolates. Samples from an individual plant were therefore assumed to contain a pool of potentially multiple genotypes of the fungus. We aligned samples to the draft *P. novopanici* genome (Gill et al. 2019) using *BWA* and deduplicated alignments with *sambamba* (Li and Durbin 2010; Tarasov et al. 2015). Since the draft genome is highly fragmented (11,088 contigs, N50=13,091bp), we performed a cleanup step by aligning contigs to a better-quality *Puccinia triticina* reference genome (Wu et al. 2021). We used relaxed alignment parameters in BLAST (-evalue 1e-5), then discarded 1254 contigs that failed to map to the *P. triticina* genome. Though a full revision of the *P. novopanici* genome is beyond the scope of this study, we used approximate contig positions in the *P. triticina* genome for visualizations, and include estimated contig scaffolding in the supplemental material. We used two pipelines to call SNPs and estimate regional diversity. For SNP-calling, we used *BCFtools* to generate an invariant-sites VCF, then filtered to biallelic sites with a minimum minor allele frequency of 0.05 that were genotyped in all of our 88 samples using *Plink2* (Chang et al. 2015; Li et al. 2009). We used *popoolation* (Kofler et al. 2011) to assess diversity between regional pools. To group reads, we used *SAMtools* mpileup (H. Li et al. 2009). We then used *Variance-sliding.pl* in *popoolation* to assess allelic diversity (π) using a sliding-window approach. We used 1000bp windows with a 100bp step size. We filtered out reads with less than 4x or greater than 70x coverage depth or an average Phred quality score less than 20. We tested for differences in pi between regions using the studentized bootstrap method suggested by Efron and Tibshirani (Efron and Tibshirani 1994). We calculated the mean π values across windows for each contig to compare between regions. We counted the number of contigs with π > 0.05 to assess the number of distinct outlier loci in each pool. We estimated F_ST_ between regions using the *grenedalf* package (Czech, Spence, and Expósito-Alonso 2023). Here we used identical filtering steps, but used individual contigs as windows to improve accuracy, then computed the mean F_ST_ per window adjusted by the number of SNPs on each contig. We performed principal component analysis using singular value decomposition through the big_SVD function in the R package *bigsnpr* (Privé et al. 2018).

### Rust phenotyping and distribution

For rust severity, we followed a similar phenotyping protocol to that previously described in (VanWallendael et al. 2020). Briefly, plants at each site were checked daily for rust presence following spring green-up. After the first instance of rust was detected, plants were scored every two weeks for at least four rating periods (8 weeks), which was typically sufficient to capture the exponential growth phase of rust infection increase. Rust was scored on a 0-10 scale, with each point corresponding to approximately 10% of the total plant leaf area covered in rust sori. We calculated the area under the disease progression curve (AUDPC) for each plant for the 8-week period centered on the inflection point of rust increase at each site. Using AUDPC rather than max or mean rust scores corrects for some of the differences in scoring rust between sites, and encompasses more temporal variation in scores. Additional phenotypic measurements were taken for switchgrass plants as described in (Lovell et al. 2021), including height, tiller number, flowering time, and biomass. We measured height as distance in cm from the base of the plant to the height of the flag leaf. We assessed flowering time as the day of year (Julian day) at which 50% of tillers were flowering. Finally, we estimated dry biomass by measuring the wet biomass of each whole plant in the field, then multiplying by moisture percent of each sample as assessed by drying a representative subsample of each plant.

#### Rust distribution and local adaptation

We assessed rust distribution across genotype, space, and time using linear mixed models in the R package sommer (Covarrubias-Pazaran 2016). Since a model including all sites failed to converge, we split sites into northern (BRKG, KBSM, FRMI, LINC) and southern (CLMB, OVTN, TMPL, PKLE, KING) regions. We used the mmer function to solve the following model with a compound symmetry variance-covariance matrix, splitting northern and southern sites. Our full model included terms for site, year, genotype, and a genotype-by-environment interaction. We tested the importance of model components by sequentially dropping each term and testing for fit differences using a likelihood ratio test via the *anova* function in *sommer*.

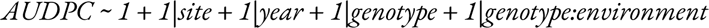

We examined relationships between rust severity and other switchgrass phenotypes using principal components analysis (PCA). We examined three focal sites, (KBSM, CLMB, and PKLE) that had the greatest number of shared genotypes (n = 1070). We used the mean value of each trait for each genotype across sites and years in the PCA. We removed 94 genotypes with missing data, then computed the PCA using the centered and scaled values in the prcomp function in R.

#### The host reciprocal transplant

We used a modified version of the classic local adaptation test to assess parasite local adaptation, which we have termed a host reciprocal transplant (Figure 2). There are several advantages in both practicality and experimental robustness to this method, though some drawbacks exist. As mentioned before, fewer regulatory and ethical issues surround transplant of hosts than their parasites, and unculturable parasites may be particularly challenging. A host reciprocal transplant overcomes some of the challenges of translating controlled common garden experiments to the field. Namely, growth chambers typically remove much of the environmental context of the host-parasite interaction, and both susceptibility and infectivity can be context-dependent. Similarly, controlled experiments select for, and are biased by genotypes that are simply better-adapted to lab or greenhouse conditions (Kawecki & Ebert 2004). However, abiotic context is quite different between parasite and host reciprocal transplants. Is local abiotic context more important for the parasite infectivity or the host susceptibility? If we make a general assumption that both host and parasite are well-adapted to their native abiotic environment, transplanting each outside their adapted range should tend to weaken populations, making hosts more susceptible and parasites less infective. Therefore, in a host transplant experiment, we would tend to underestimate the strength of local adaptation, since foreign parasite populations infect hosts with lowered susceptibility. In contrast, parasite transplants may overestimate local adaptation, since foreign parasites may have reduced infectivity in novel environments. A reversed pattern may be possible in systems where organisms are maladapted to their abiotic environment, or if interactions with other organisms such as hyperparasites make foreign transplant advantageous.

**Figure 2:**
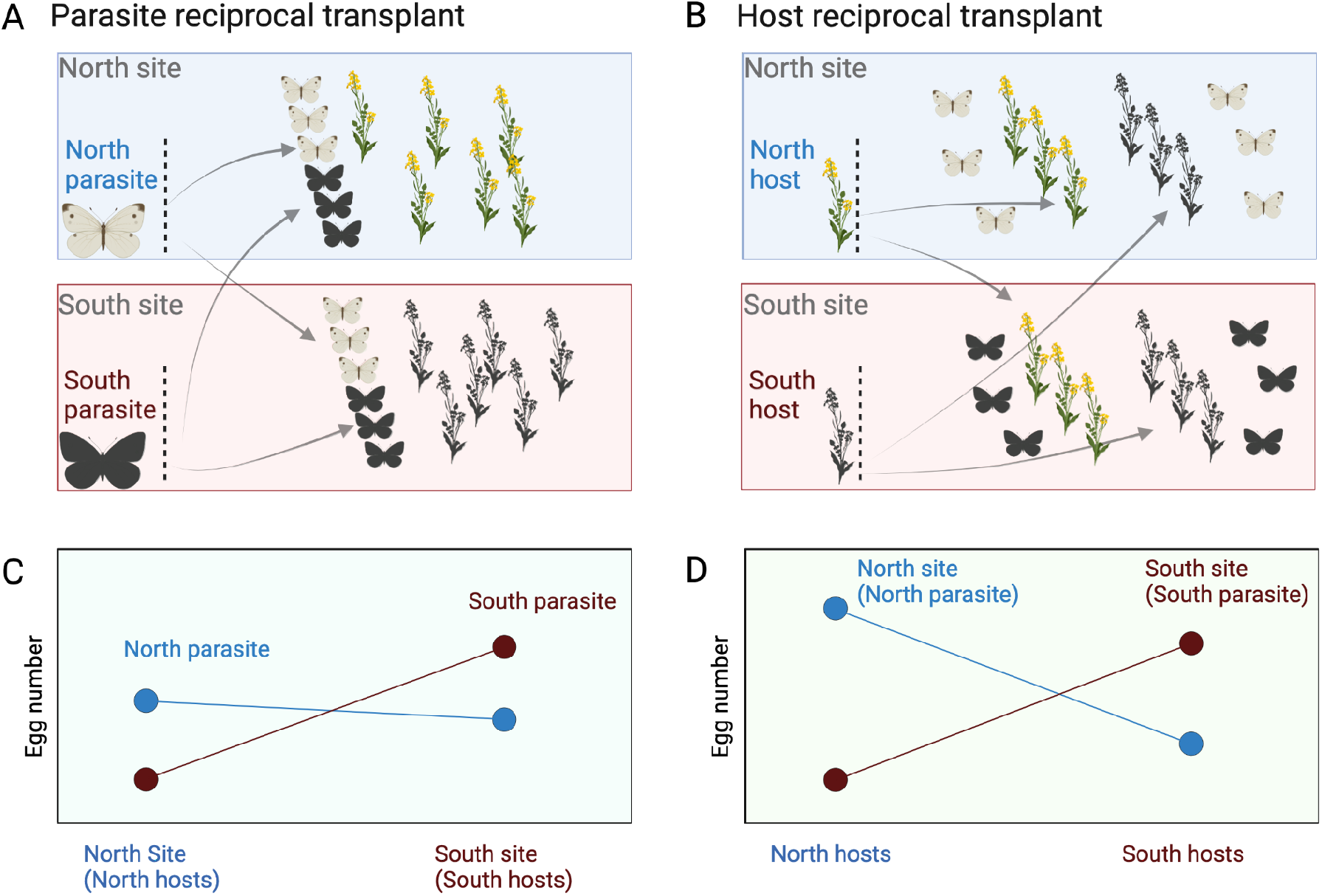
Comparing two methods for testing parasite local adaptation, a parasite reciprocal transplant (AC) and a host reciprocal transplant (BD). In this example scenario, two host populations (field mustard plant icons) and two parasite populations (cabbage white butterfly icons) are differentiated between two sites, North and South. **A** traditional parasite reciprocal would introduce northern and southern butterflies into mustard populations at each site. **B.** A host reciprocal transplant would introduce northern and southern mustards in a common garden to butterfly populations at each site. **C.** Proof of local adaptation via the local-foreign comparison would require greater fitness for north butterflies over south butterflies on north mustards, and greater fitness for south butterflies over north butterflies on south mustards. This is indicated by more eggs produced by the north parasite (blue points) in the north site, but more eggs produced by the south parasite at the south site (red points). **D.** Proof of local adaptation would require the same fitness advantage as in a parasite transplant: north butterflies over south butterflies on north mustards, and greater fitness for south butterflies over north butterflies on south mustards. However, this test would be between sites rather than within sites. In the plot, the x-axis indicates host population, while the color of points indicates parasite population at each site.

Ultimately the utility of parasite or host reciprocal transplants depend on the specifics of the study system. Several potential pitfalls are possible when using this method. If parasite populations are panmictic, or ranges change rapidly across years, they are unlikely to be locally adapted, and will be harder to assess. Transplanting a host that contains potential parasite propagules will confound the results of the experiment, (though post-hoc testing of diseased tissue to assess parasite source can alleviate this). Therefore, extra care must be taken to remove vertically transmitted parasites from transplanted hosts. This study used a systemic fungicide before transplantation. Finally, site distance should be adjusted according to parasite dispersal capacity. If the same parasite source population infects multiple sites, results may be biased. While long-distance dispersal is possible in rusts, most infection occurs from locally dispersed spores (Farber, Medlock, and Mundt 2017).

To specifically test the hypothesis that rust is locally adapted to switchgrass, we assessed whether rust BLUPs were higher in genotypes facing coevolved pathogens. Since BLUPs were not normally distributed (Shapiro-Wilk test p < 0.0001), we used a nonparametric Dunn’s test to assess differences in means between groups, with a Bonferroni correction for multiple testing. We performed an additional simple test for switchgrass local adaptation to rust by regressing distance between planting sites with collection locations (Figure 1) with rust severity AUDPC using a linear model. This test is not diagnostic of local adaptation, but is an indicator of the degree to which the match between native environmental conditions matters for rust infections.

### GWAS data analysis

In our analyses we used a set of SNPs first generated by Lovell et al. (Lovell et al. 2021). This set comprised 10.8 million SNPs after quality filters and a minor allele frequency cutoff of 0.05. We performed a PCA using the *big_SVD* command in the R package *bigstatsr* (Privé et al. 2018). We computed GWAS using the *pvdiv_gwas* function in the *switchgrassGWAS* R package (Lovell et al. 2021). This function runs linear regression on filebacked big matrices using the *big_univLinReg* function in *bigstatsr*, with ten principal components as covariates to correct for population structure.

A major challenge in this study lay in correcting for heterogeneity across time and space. Since rust pathogen pressure can be variable across years and different technicians rated rust across sites using a relatively subjective metric, finding the true genetic basis of rust resistance required minimizing other sources of variation. When combining data across environments, genetic mapping studies often calculate best linear unbiased predictors (BLUPs) for each genotype, then use these as a phenotype for GWAS or QTL (quantitative trait locus) mapping (Wallace et al. 2016; Cui et al. 2021; Kumar et al. 2018). BLUPs estimate the random effect of genotype in multi-environment linear mixed models, with the goal of distinguishing genetic from environmental effects. We used the above described LMM to calculate

BLUPs for northern and southern regions. BLUPs correlate well with the mean of centered and scaled AUDPC genotype scores for both northern and southern sites (r = 0.87 and r = 0.94, respectively). Using a kinship matrix instead of genotype identity in the model produced highly similar BLUPs (r = 0.99 for North and South), so we only used genotype identity.

We considered genes <10Kb from an outlier SNP as linked to the SNP, since this is the typical distance of linkage decay in the switchgrass genome. We investigated gene function and homology using JGI JBrowse (Skinner et al. 2009), NCBI BLAST, and NCBI Genbank. We assessed which gene functions were overrepresented in our dataset using a bootstrapping method focused on the 20 gene functional annotations with the greatest difference in frequency between the sample and total dataset. We randomly permuted genes from the total genome-wide list for 100000 iterations, then assessed whether each function was more common than the permuted set <5% of the time.

To provide independent lines of evidence towards candidate genes, we used data from a previously described RNA-sequencing experiment (VanWallendael et al. 2022). In short, this experiment used transcript abundance from the uppermost fully expanded leaves from a single tiller of each genotype at phenologically similar dates at each site: PKLE: 8-June 2016; CLMB: 21-July 2016; KBSM: 18-Aug 2016. RNA was assayed from leaf tissue that was (1) excised at the ligule, (2) 2 cm of the proximal portion of the excised leaf were separated from the midrib, (3) placed in a 2 mL Eppendorf tube loaded with three stainless steel beads, (4) immediately frozen in liquid nitrogen and (5) transported on dry ice to the laboratory. Tissue was homogenized with a Geno/Grinder 2000. RNA was extracted with the standard Trizol protocol and treated with DNase I to remove contaminating genomic DNA. We compared leaf RNA expression in four genotypes at three of our study sites, KBSM, CLMB, and PKLE. These four genotypes contained two that are typically rust-susceptible (DAC6 and VS16) from the upland ecotype, and two that are more resistant (AP13 and WBC3) from the lowland ecotype. We assessed differential expression at focal genes using the *DeSeq2* package in R (Love, Anders, and Huber 2014). We filtered out genes that had counts lower than 10 for three or more samples, according to best practices (Love, Anders, and Huber 2014).

Finally, we compared our results to outlier regions for other traits identified in Lovell et al. (2021). This study used Bayesian tests to combine data across multiple environmental phenotypes. We checked for concordance between outliers by checking the proportion of hits within overlapping 10Kb windows using custom scripts. Since many of these SNPs are clustered, we first collapsed 5504 outliers into multi-SNP loci. In 100000 permutations, we randomized the genomic position of each locus, then counted the number of overlapping loci with non-rust traits. In addition, we assessed the genetic correlation of rust to other switchgrass phenotypes using mixed models in R. We fit three models for biomass, flowering time, and tiller count that included environment (site-year combination) and a kinship matrix with the formula:

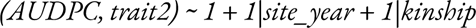

Biomass and tiller count were taken at the end of the season, and represent the dry mass of aboveground leaf tissue, and the number of tillers (stems) respectively (Lovell et al. 2021). Flowering time was assessed as the day of year when half the crown had flowered.

## Results

### Rust population genetics

We examined rust populations on leaves in multiple locations with the goal of determining how these differ genetically using whole-genome resequencing. To assess population structure, we called SNPs in 88 samples across nine sites by aligning reads to the *P. novopanici* genome (Gill et al. 2019). This resulted in a set of 2.4 million SNPs with a minimum MAF (minor allele frequency) of 0.05 that were called in all samples. Since rust isolated from a single plant may include multiple genotypes, individual isolates are treated as pooled samples. We performed PCA using single vector decomposition, and found that within shared SNPs, samples from northern and southern sites were mostly distinct (Figure S1) and that PC1 was correlated with latitude of collection (Figure 3A).

**Figure 3:**
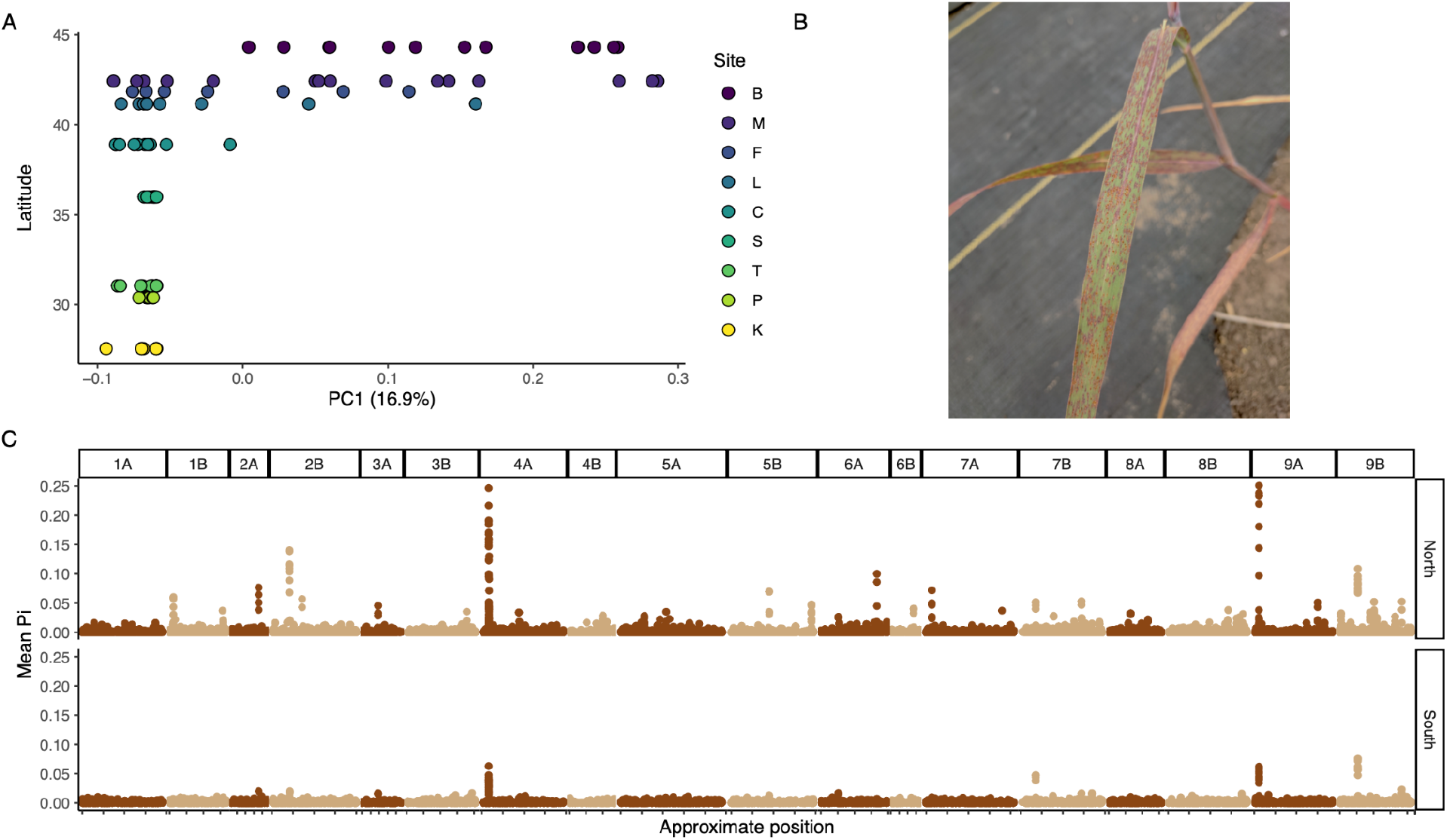
*Puccinia novopanici* leaf rust population genetics across nine sites. **A.** The y-axis indicates latitude of the collection site, and the x-axis is the first principal component of a PCA of 2.4 million SNPs. Site codes correspond to: B-Brookings, SD; M-Kellogg Biological Station, MI; F-Fermilab, IL; L-Lincoln, NE; C-Columbia, MO; S-Stillwater, OK; T-Temple, TX; P-J.J. Pickle Research Campus, TX; and K-Kingsville, TX. **B.** *P. novopanici* sori under field conditions in Kingsville, TX. Photo by Acer VanWallendael. **C.** Mean nucleotide diversity (π) across 10kb windows in the *P. novopanici* genome. Positions are shown by mapping location in the *P. triticina* genome.

In order to uncover differences in genetic diversity between rust populations, we scanned the genome for regions of increased genetic diversity using pooled sequence analyses. In this analysis, individual and site differences are combined to highlight larger-scale differentiation across the genome. Since rust dispersal within a season occurs mostly through asexual urediniospores, we expected to find mostly low-diversity clones. However, high-diversity windows may indicate emergence of clones with novel mutations in fitness-related genes. We examined northern and southern regional pools (Figure 3C) for loci with increased diversity that may indicate selection. Since the *P. novopanici* genome is highly fragmented, we anchored contigs to the haplotype-resolved dikaryotic *Puccinia triticina* genome for organization. One large outlier locus with π > 0.75 on Chromosome 9B was linked to the Internal Transcribed Spacer (ITS) region. Since ITS regions often exist as large tandem repeats, the very high diversity at this locus is likely spurious, so we excluded this region from further analysis. Overall genome-wide π was 4.7% higher in the North (northern π = 7.97 × 10^-4^, southern π = 7.61 × 10^-4^, bootstrap p = 0.0006). Additionally, northern populations contained 15 unique outlier loci (π > 0.05; Figure 3C), whereas the southern populations contained just three outliers. Genome-wide average F_ST_ between regions was moderately high at 0.155 (range: Chr 4B = 0.131 - Chr 8A = 0.165). While gene annotation exists for some *Puccinia spp.*, functional annotation is limited to a few well-conserved genes. Therefore, we were not able to confidently link outliers to known rust genes.

### Variation in Rust Susceptibility

The GWAS panel in this study represents one of the most extensive multiple common garden studies of a perennial plant created to date. We used the diverse source material planted at multiple locations to measure both variation in the genotypic component (source genotype), and environmental component (year and planting site) of the expression of rust susceptibility. In order to isolate genetic contributions to rust susceptibility, we aimed first to estimate variation in the environmental component of rust severity.

Since a global linear mixed model (LMM) for all sites and years failed to converge, we conducted most analyses with northern and southern sites separated. To compare the relative amounts of variation attributable to genotype, site, year, and Genotype x Environment interaction (GxE) we fit an LMM for AUDPC across sites and years. For both regions, the highest variance components were genotype (northern = 0.244, southern = 0.297; Table 1), followed by GxE (northern = 0.264, southern = 0.109). Overall, the model explains 63.2% of the variation in AUDPC in northern sites (AIC = 3788), and 55.5% of variation in southern sites (AIC = 2550). Dropping each of the terms from models resulted in significantly poorer model fit (Likelihood Ratio Test; Table 1).

**Table 1:** Variance proportions for LMMs in northern and southern regions for rust severity. LRT columns show the results of a Likelihood Ratio Test that compares a full model and a model without each term. The GxE term represents a genotype-by-environment interaction, with each site-year combination as a separate environment.

| **Northern** |  |  |  |  |
| --- | --- | --- | --- | --- |
|  | **Prop. variance** | **Zratio** | **LRT ChiSq** | **LRT pval** |
| Site | 0.0772 | 1.21 | 432.8 | < 0.001 |
| Year | 0.0472 | 0.99 | 220.6 | < 0.001 |
| Genotype | 0.2439 | 12.75 | 621.4 | < 0.001 |
| GxE | 0.2635 | 6.67 | 16.2 | < 0.001 |
| *Residual* | 0.3682 | 9.62 |  |  |
|  | **LogLik** | **AIC** | **BIC** |  |
|  | -1893 | 3788 | 3794 |  |

| **Southern** |  |  |  |  |
| --- | --- | --- | --- | --- |
|  | **Prop. variance** | **Zratio** | **LRT ChiSq** | **LRT pval** |
| Site | 0.107 | 0.99 | 327.9 | < 0.001 |
| Year | 0.042 | 0.99 | 172.2 | < 0.001 |
| Genotype | 0.297 | 12.93 | 891.0 | < 0.001 |
| GxE | 0.109 | 2.49 | 5.7 | 0.017 |
| *Residual* | 0.445 | 10.17 |  |  |
|  | **LogLik** | **AIC** | **BIC** |  |
|  | -1274 | 2550 | 2572 |  |

Different technicians rated rust severity at each site, so the variance explained by site may be partially attributable to subjective rating differences (Figure 4A). Some sites had consistently high rust ratings, such as CLMB (Columbia, MO), and others varied between years like in BRKG (Brookings, SD), which had no rust in 2020 or 2021. To reduce this bias when summarizing across genotypes, we centered and scaled rust severity scores before comparing populations (Figure 4B). Rust varied greatly between genetic subpopulations (Figure 4B), and covaried with several other traits, particularly time of plant green-up and biomass (Figure 4C). The Atlantic population had the lowest overall rust severity and the Midwest had the highest. The Gulf population was split between genotypes originating from the Gulf coast and those from more inland, mostly in Texas, so these subgroups are separated in most analyses. Octoploid genotypes, which can be found across the switchgrass range, had the greatest overall variation. In a trait PCA (Figure 4C), biomass clearly was negatively correlated with rust scores, with higher biomass present in the central Texas population, and higher rust scores in the Midwest. Greenup date was more closely correlated with rust score, and traded off with tiller count.

**Figure 4:**
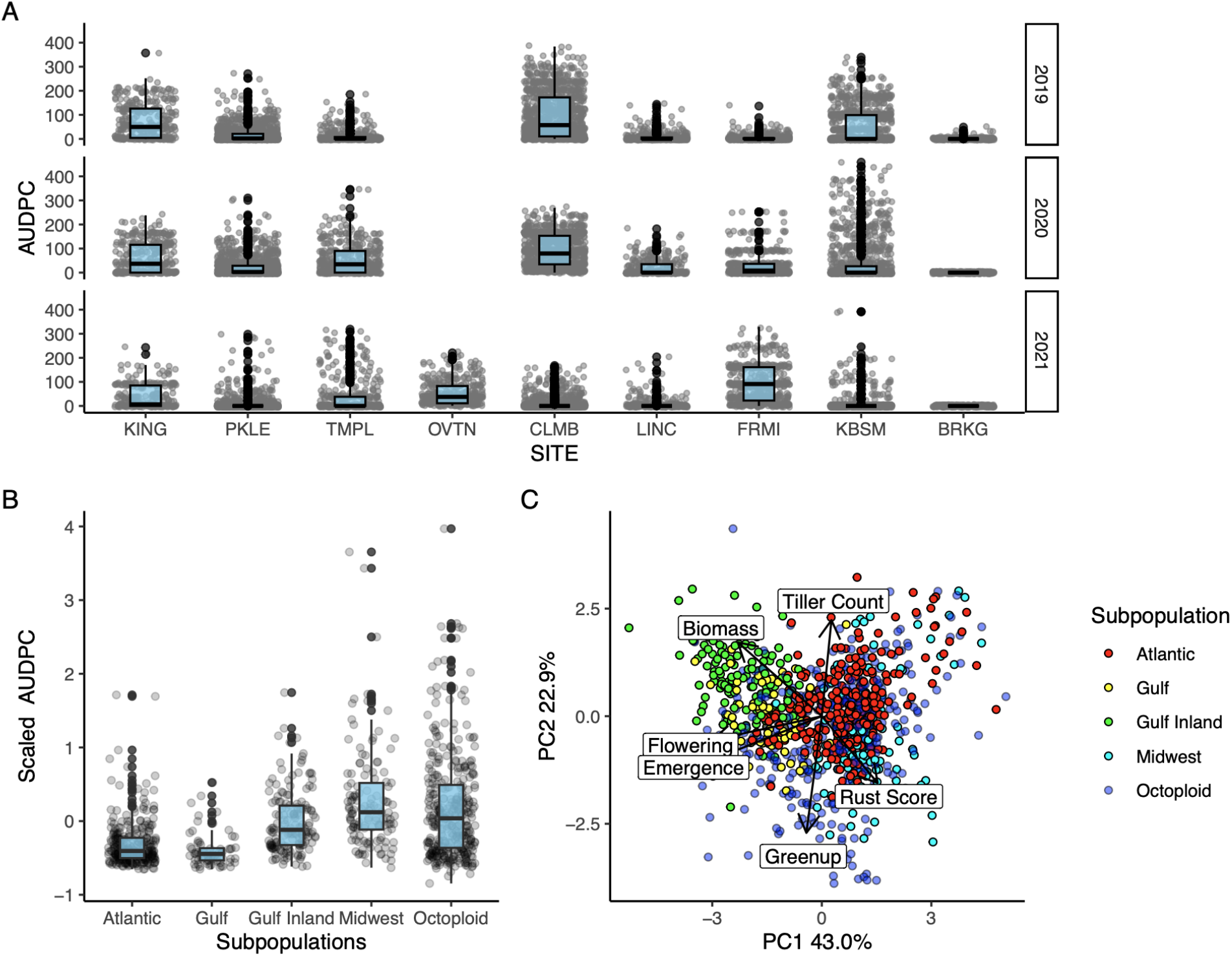
Rust score variation across populations, space, and time. **A.** Site & year variation in rust infection. Points shown indicate the area under the disease progression score across 8 weeks for each plant in each year. Sites on the x-axis are shown from southmost (KING: Kingsville, TX) to northmost (BRKG: Brookings, SD). 2019-20 data could not be collected at OVTN (Overton, TX) and rust was not found in BRKG in 2020-21. **B.** Genetic populations vary in rust severity. Points indicate mean scaled rust severity (AUDPC) across sites and years. The Gulf genetic population has a subdivision between genotypes originating inland and those by the coast. **C.** Phenotypic PCA biplot for major traits across all sites and years. Points are colored by subpopulation.

To test the specific hypothesis that rust pathogens are locally adapted to switchgrass genotypes, we examined differences in rust severity for genotypes across locations. To separate environmental from genetic effects on rust severity, we calculated Best Linear Unbiased Predictors (BLUPs) from the genotype term of the previously mentioned linear mixed-effect models. BLUPs predict genotypic contribution to rust severity, with higher values indicating genotypes with greater susceptibility to rust across sites and years. Individuals from the Midwest population were commonly susceptible to rust present in northern sites (positive BLUP; Figure 5A), whereas those in the Gulf inland subpopulation were more susceptible when transplanted to southern sites (Figure 5B). Gulf and Midwest populations therefore conformed to expectations of local adaptation, with higher rust severity in southern and northern planting sites, respectively (Figure 5C; Dunn’s test: Bonferroni-adjusted p < 0.0001 for both populations). In contrast, rust severity was not differentiated in Atlantic genotypes across regions (Figure 5C; Dunn’s test: Bonferroni-adjusted p > 0.999).

**Figure 5.**
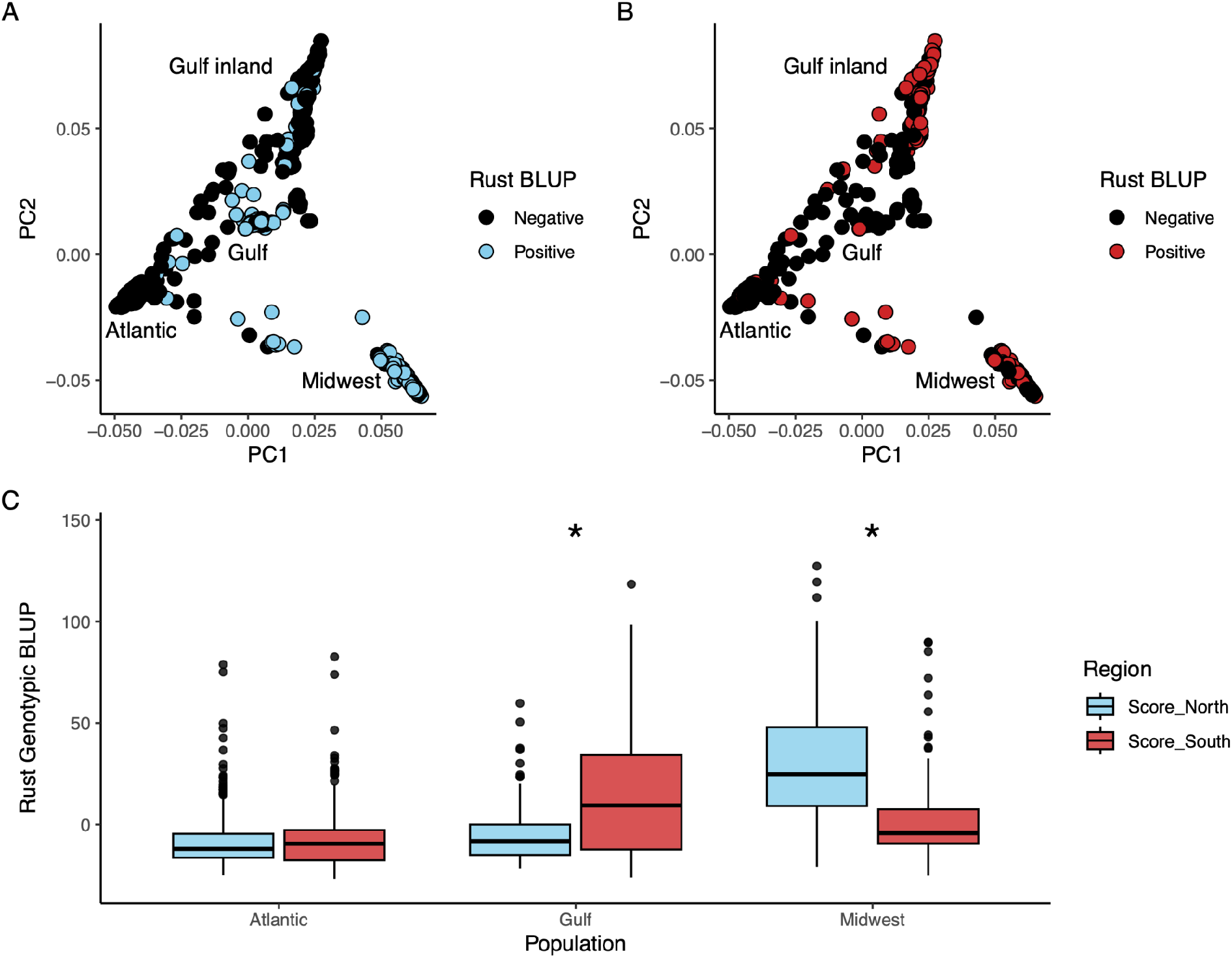
Switchgrass genomic principal component analysis showing susceptibility to endemic rust in northern (**A**) and southern (**B**) sites. Rust severity BLUP was measured across northern and southern sites. Negative scores indicating genotype resistance are shown as black, positive scores indicating genotype susceptibility are colored. **C.** Rust BLUPs (susceptibility) by host group and region. Blue boxes indicate samples planted in northern common gardens, red boxes indicate samples planted in the southern gardens. Asterisks indicate regional differences; p < 0.0001 for both.

To test the impact of environmental differences in common garden locations on switchgrass susceptibility to rust, we examined the relationship between distance from location of origin (source population for each genotype) and rust infection at each growing site. To minimize the large confounding effects of subpopulation, we conducted separate tests within each population, then used a multiple-testing correction. The Midwest and Octoploid populations showed a locally maladaptive pattern, with a significant negative relationship between distance from location of origin and rust infection (Table 2). For the Midwest population, this relationship resulted in an average of 1.90% lower rust severity for every 100 kilometers from the origin. The Atlantic population, conversely, showed a positive relationship between distance from location of origin and rust severity, while the Gulf population showed no relationship. For all populations, however, this relationship explained relatively little (< 6%) of the variation in rust severity.

**Table 2:** Testing for switchgrass local adaptation to rust populations. Linear model for the relationship between distance from location of origin and growing location and rust severity.

| Pop | Coeff | tstat | p | R2 | n |
| --- | --- | --- | --- | --- | --- |
| **Midwest** | **-3e-05** | **-3.080** | **0.00224** | **0.027** | **338** |
| Gulf | -1e-05 | -1.644 | 0.10075 | 0.004 | 609 |
| **Atlantic** | **2e-05** | **4.945** | **0.00000** | **0.032** | **742** |
| **Octoploid** | **-4e-05** | **-5.974** | **0.00000** | **0.054** | **629** |

### GWAS

We conducted GWAS separately for the northern and southern regions, using rust severity BLUPs as predicted traits in a linear GWAS model. We found numerous loci associated with rust severity, indicating a largely polygenic response (Figure 6). In both northern and southern regions, p-values showed signs of inflation, despite a conservative correction with 10 PCs (North lambda = 1.042, South lambda =1.092; Figure S2). For all significant SNPs shared between regions, there was only a weak positive correlation between -log10 p-values (r = 0.354, p < 0.0001), indicating few shared mechanisms. However, the correlation disappears when examining only the top SNPs in each region. Curiously, the average effect sign (which indicates whether the major or minor allele had a higher trait value) of top SNPs was also different between regions (Figure S3). While top SNPs at northern sites tended to have lower BLUPs with the minor allele, top SNPs at southern sites had lower BLUP values with the major allele. This may indicate that when genotypes are grown in the south, alleles that are relatively common in the diversity panel contribute the most to resistance, whereas in the north resistance comes from rarer alleles.

**Figure 6.**
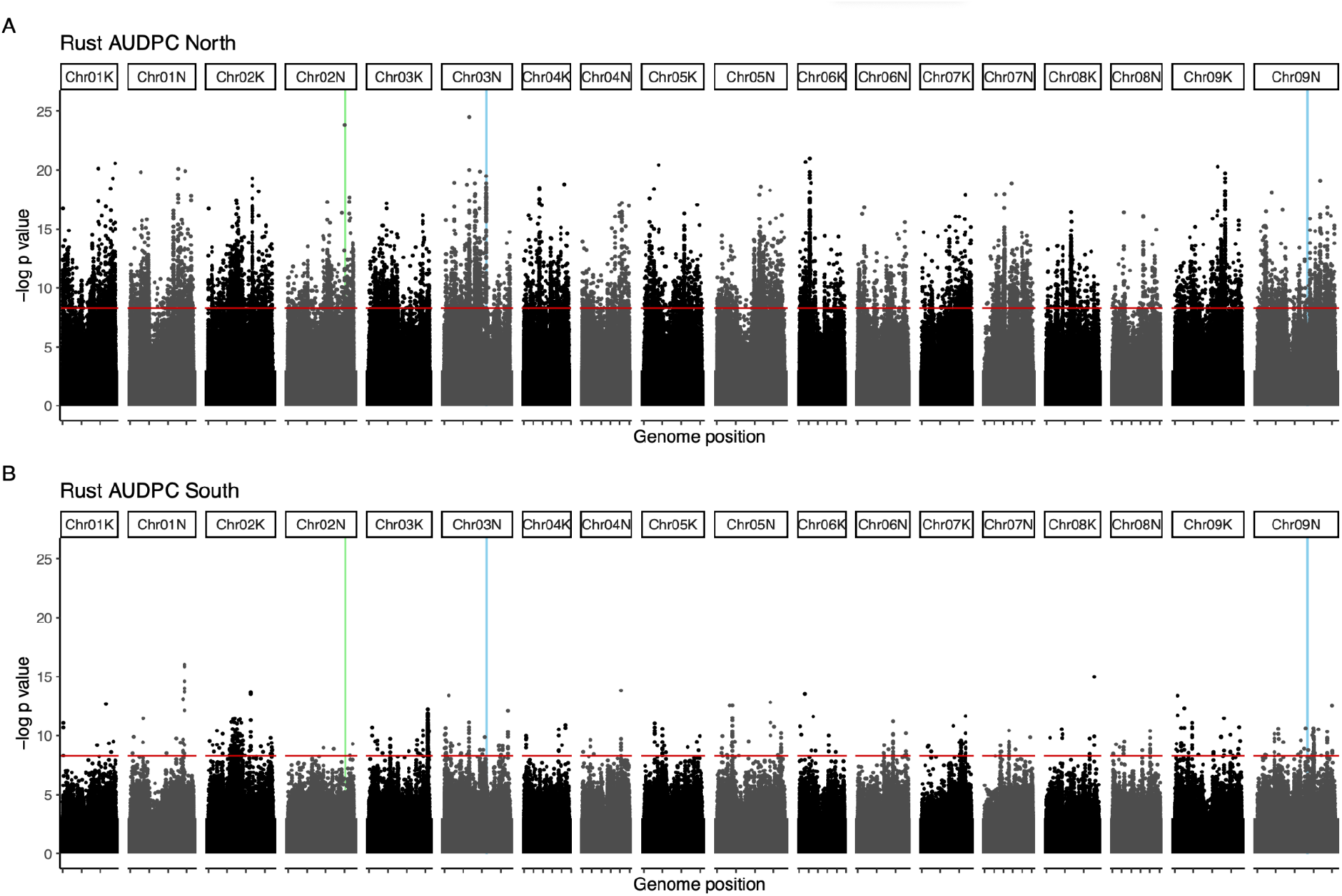
GWAS of mean scaled AUDPC in northern (**A**) and southern (**B**) regions. Vertical lines indicate the positions of candidate loci found in previous experiments (VanWallendael et al. 2020; VanWallendael et al. 2022). The green line on chromosome 2N indicates a microbiome structure - associated GWAS outlier, and blue lines indicate rust resistance QTL *Prr1* (Chr03N) and *Prr2* (Chr09N). Red horizontal lines indicate a Bonferroni cutoff.

We examined genes linked to the top 100 outlier SNPs in the northern and southern sites. There were no overlaps between top SNPs or the genes linked to them. We used enrichment analysis to determine which gene annotations were overrepresented in genes linked to top SNPs. While most of the enriched annotations did not have a clear link to disease resistance, there were some exceptions. In the north, four gene annotations were overrepresented in our top SNPs (Figure S4). Terpenoid cyclases, which allow formation of specialized metabolite defenses such as antifungal leaf saponins, were common (p < 0.001). In the south, the most notable of the five overrepresented annotations were EF-TU receptors, Leucine-rich repeat receptors (LRRs) that typically recognize bacterial pathogens (Schoonbeek et al. 2015). However, LRR proteins have highly variable structures and may act as receptors for a wide range of PAMPs (pathogen-associated molecular patterns), so these may be fungal rather than bacterial receptors. A surprising result was the overall difference in frequency of LRRs across regions. Though these genes are common across the genome, few were linked to top SNPs in the north, while several were linked in the south. Outliers on chromosomes 2N (North), 3K (South), 4K (South), and 6K (South) were closely linked to LRRs. A MATE (multidrug and toxin extrusion) transporter is linked to an outlier on chromosome 1K in the south. These genes can function in pathogen responses by moving hormones such as salicylic acid in response to infection (Serrano et al. 2013).

Previous research identified several loci associated with leaf fungi (Milano et al. 2016; VanWallendael et al. 2020; VanWallendael et al. 2022). The strongest candidate gene we identified within the chromosome 3N *Prr1* QTL region was Pavir.3NG168388 (Figure S5). This gene is an oligopeptide transporter similar to YELLOW STRIPE-LIKE (YSL3), which functions in metal ion distribution in plants (Sheng et al. 2021), but has a notable role in plant-pathogen interactions (Chen et al. 2014). The *Prr2* QTL region on 9N contained relatively weaker GWAS hits, but one of these was linked to the highly conserved CCR4-NOT complex (Pavir.9NG474500), which acts as a multifunctional gene expression regulator (Collart and Panasenko 2017). A large outlier on chromosome 2N is near a previously-discovered cluster of cysteine-rich receptor-like kinases associated with variation in the leaf microbiome (VanWallendael et al. 2022). This outlier is linked more closely, however, to four genes with predicted functions in drug- and disease-resistance (Pavir.2NG514900.1, 515000.1, 515100.1, and 515300.1; Figure S5). Homeologs of three of the four genes are also found on chromosome 2K, though there is no associated outlier SNP on this chromosome.

The large outlier region on chromosome 2K in the south encompasses nearly the entire pericentromeric region of this chromosome, which has a highly suppressed rate of meiotic recombination (Lovell et al. 2021). The pericentromeric region is also gene-poor, but contains some potentially relevant genes. In particular, one gene encodes a NOI4-like protein, which is within the cascade of immune responses to effectors in Arabidopsis (Redditt et al. 2019). This locus also contains an *FCA* (flowering time control) ortholog, as well as several genes involved in DNA replication and repair such as DNA helicase and gyrase, as well as *SMG7*, an essential gene in meiosis (Riehs et al. 2008).

We collected RNA-seq data from switchgrass leaves in three of our sites, KBS, Hickory Corners, MI; Columbia, MO; and JJ Pickle Research Station, Austin, TX. To better explore causal loci, we analyzed transcript abundance of genes underlying GWAS peaks across four genotypes, two upland cultivars, and two that are lowlands. Of the 107 genes closely linked to top GWAS hits in the North, 82 were differentially expressed between resistant and susceptible varieties (top 20 in Table S1). A gene with unknown function (Pavir.7NG105945) was most clearly differentiated, followed by a gene on 1N in the RING/U-box superfamily. The oligopeptide transporter underlying the *Prr1* QTL had very low expression in the leaf for the four genotypes in this study, but the gene cluster on chromosome 2N was differentially expressed. Three terpenoid cyclases on chromosome 1K were minimally expressed in southern lowland genotypes, particularly when grown in the northernmost site (Figure 7). For southern outliers, 96 of 130 genes linked to top GWAS hits were differentially expressed. The strongest differentiation was seen in a small ribonucleoprotein F gene on chromosome 5N (Table S1), but a Leucine-rich repeat kinase detected on chromosome 3K (Pavir.3KG551700) was clearly differentiated as well.

**Figure 7:**
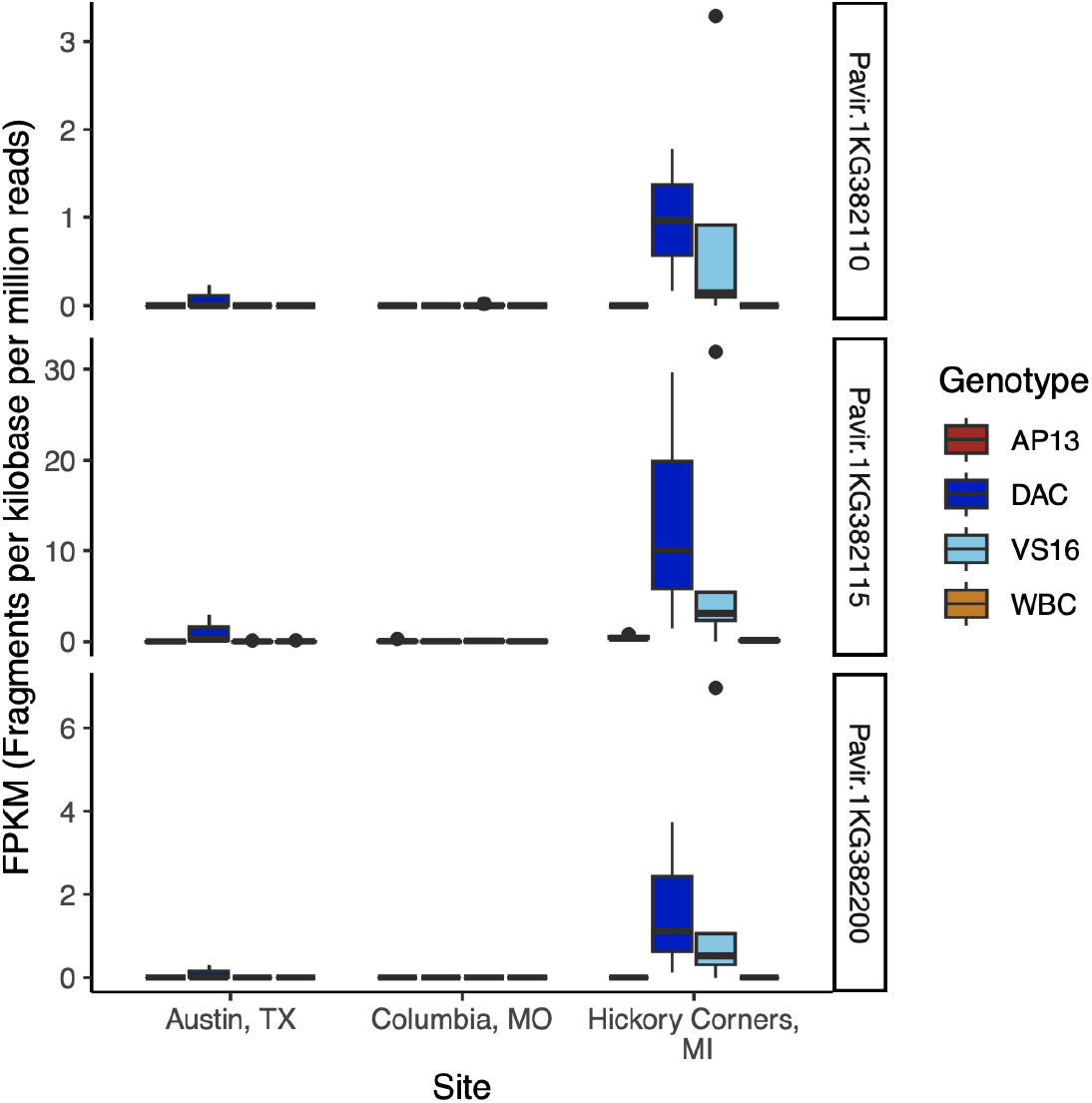
Expression differences for three terpenoid cyclase-encoding genes linked to large GWAS outliers on chromosome 1K. Leaf tissues from four genotypes were tested at three sites in 2016. AP13 & WBC are southern genotypes in the Gulf population; DAC & VS16 are northern genotypes in the Midwest population.

### Links to other traits

To determine the extent to which variation in rust severity is linked to other switchgrass phenotypes, we assessed genetic correlations with biomass, flowering time, and tiller count using LMMs. As was suggested in a trait PCA, all three traits had negative genetic correlations with rust severity. Flowering time had the strongest correlation (-0.410) followed by tiller count (-0.342), and biomass (-0.254). To understand the physical relationship between traits, we examined overlap between 11834 GWAS outliers for rust severity with 12231 GWAS loci for other traits identified in Lovell et al. (Lovell et al. 2021), including biomass, survival, and climate-associations. We merged GWAS outliers into 3236 rust-related loci and 8643 biomass, survival, and climate loci using 20kb windows around each SNP to define linkage, since that is the typical inflection point at which linkage disequilibrium decay flattens (Lovell et al. 2021). In both northern and southern GWAS, about half of rust loci (North = 1486/2961, South = 141/275) overlapped with climate, biomass, and survival loci.

To test if this number of overlaps could be attributable to chance we used bootstrapping to permute the locations of rust-related loci throughout the genome. In 100,000 permutations, we did not observe any cases where the number of overlapping loci was greater than the observed numbers in northern GWAS (p < 0.00001), and just two cases in southern GWAS loci (p = 0.00002), so we can infer that there was significant concordance between rust GWAS outlier regions and those for other switchgrass traits (Figure S6). Overlapping loci were not strongly enriched for either fitness measured by end of season biomass- or climate-related traits in the North (1.649% fitness related traits in the overlapping set versus 1.473% in the original set). However in the South, only one rust locus overlapped with a biomass-related locus (0.709%). The loci with the highest scores for both rust severity (-log(p) > 10) and other traits (bf > 10) were on chromosomes 1N, 2K, 3K, and 3N. These are linked to genes involved in cellular signaling pathways, including a MYB gene on chromosome 1N and SEC14-like genes on both chromosomes 2K and 3N (Table S2).

## Discussion

Predicting whether a host or parasite will be more successful in a coevolutionary arms race is a challenge that requires understanding the genetic makeup of both organisms, as well as the environmental context. Theory holds that parasites with short generation times will locally adapt across the geographic range of the host more quickly than the host can evolve defenses (Gandon and Michalakis 2002). However, few systems have been studied under natural conditions, and little is known about how host genetic associations can differ between locally adapted parasite populations. In this study we followed natural rust infection cycles in switchgrass planted at multiple locations over three years to understand the genetic and geographic structure of the pathosystem. We confirmed our predictions that rust would be genetically differentiated across northern and southern regions, and found that it was broadly locally adapted to switchgrass genotypes sourced from those regions using a version of a reciprocal transplant experiment. Further, we found that earlier results using QTL mapping were recapitulated with GWAS, showing different patterns of genetic association in northern and southern sites, and confirming the importance of a QTL on chromosome 3N. The top candidate for this QTL is Pavir.3NG168388, a YSL3 metal ion transporter ortholog. The higher resolution of GWAS allowed us to show that genes linked to top outlier SNPs were enriched for different cellular functions across regions, although there is no clear pattern in enriched functions in northern or southern field sites. Further, we found that the switchgrass genes associated to rust variation are often linked to climate-related loci, indicating that adaptation to this rust species must be studied with its regional abiotic context.

### Rust is locally adapted in northern and southern switchgrass populations

Studying local adaptation in microbes poses a unique challenge, since the scale of what constitutes “local” can vary by orders of magnitude depending on the species (Kraemer and Boynton 2017). Host-parasite systems are useful in this respect, since the scale of local adaptation is constrained to the host range. As such, a number of studies have uncovered local adaptation in host-associated microbes (reviewed in (Kraemer and Boynton 2017), including a number in rust-plant pathosystems (ex. Thrall and Burdon 2002; Laine et al. 2014; Cassetta et al. 2023). However, it is rarer to uncover a host-pathogen system wherein the host has been shown to be locally adapted to abiotic environmental conditions at a similar scale as its locally adapted pathogen. In most models of parasite local adaptation, parasites are treated as the dominant selective force on hosts, so pathogen local adaptation necessarily implies host maladaptation (Lemoine, Doligez, and Richner 2012). However, in natural systems, plants experience a mosaic of selection from both biotic and abiotic factors that may result in intermediate cases. In *Holcus lanatus*, for instance, researchers found that local plants showed greater infection from a rust fungus, but overall greater fitness than foreign transplants, probably since rust infections occurred later in the plant life cycle and therefore had a smaller relative fitness effect (Crémieux et al. 2008). In switchgrass, rust infection causes clear damage to plants, and is negatively correlated with biomass, a proxy for fitness (VanWallendael et al. 2020). However, even stronger abiotic selective events appear to play an essential role in switchgrass local adaptation. Freezing events prevent southern switchgrass genotypes from invading northern regions, causing substantial mortality in some years (Lovell et al. 2021). Since the rust impact on fitness is proportionally lower, lower rust disease load of southern switchgrass genotypes grown in the north has negligible fitness benefit compared to the strong negative selection imposed by freezing. This finding is consistent with predictions that parasites often evolve to more moderate levels of damage to hosts over time (Lenski and May 1994), and suggests that a common portrayal of host-parasite local adaptation as a zero-sum arms race is oversimplified.

Measuring rust local adaptation depends on our capacity to show that rust populations are different across the regions of interest. Historical models such as the “everything is everywhere, but the environment selects” adage assume very limited barriers to dispersal in microbial species, with local environmental conditions the main determinant of success of particular species or populations (Baas-Becking 1934). Improved methods of detection have helped to delineate the actual dispersal ranges of many species (Golan and Pringle 2017). In rust species, spore dispersal can sometimes occur over intercontinental distances (Hovmøller, Thach, and Justesen 2023), but long-distance dispersal is less likely to explain consistent seasonal variation. Environmental selection after dispersal may occur directly through rusts’ freezing susceptibility, or indirectly through range limits of susceptible hosts. While many cereal rusts have two hosts corresponding with different life stages (Ono 2002), the alternate host is not definitively known for switchgrass rust (Demers et al. 2017). Our results show a difference in rust populations corresponding approximately with 42° N latitude, or USDA Plant Hardiness Zone 6a (“2023 USDA Plant Hardiness Zone Map”). While the exact position of the division may shift according to yearly climatic variation, consistent results in common garden studies indicate that it is an important division for fungal variation. Since we previously found evidence for a North-South division in the non-rust switchgrass leaf-associated fungal microbiome (VanWallendael et al. 2022), it is more likely that rust population differences are driven by direct environmental selection, rather than through differences in primary or secondary hosts.

### The genetic basis for resistance differs across regions

While many phenotypic traits have a complex genetic architecture, with potentially hundreds of contributing alleles, disease resistance is notable since it is often attributable to one or a few large-effect genetic loci. In qualitative resistance, variation in just one or two R-genes such as NB-LRRs can completely prevent infection by a pathogen (Eitas and Dangl 2010). In this study, we saw that variation in disease severity was associated with numerous genetic loci across the genome. This pattern is more commonly associated with quantitative resistance, when numerous genes contribute to variation in disease, often without a clear link to the plant hypersensitive immune response (González, Marcel, and Niks 2012). While we saw that rust severity was associated with variation in several potential immune receptors, we found many other genes with less clear links to disease resistance. Some of these genes may be important candidates for improving quantitative resistance to switchgrass rust, but others may be artifacts of strong population structure in switchgrass. Additionally, alternative mechanisms for avoiding disease may result in a more complex trait architecture.

In addition to the complexity of the underlying genetic architecture, our ability to identify causative loci was impacted by the high level of population structure among major groups of switchgrass (Gulf, Atlantic, Midwest). Population structure is a major challenge for GWAS (Berg and Coop 2014). When traits show greater variation among than within populations, many neutral loci that differ in allele frequencies among populations can be spuriously associated with unrelated traits. In an attempt to mitigate this issue, we used a strong correction for population structure by including ten principal components of genetic variation as covariates in our model. We also used a conservative

Bonferroni-adjusted cutoff to correct for multiple testing. In contrast to our current study, our previous QTL mapping results (VanWallendael et al. 2020) do not suffer from population structure bias and thus, provide a useful benchmark for comparison. Since we recovered the two large-effect loci *Prr1* and *Prr2* in the north, and found a similar north-south divergence in their association, we are confident that many large-effect GWAS loci are likely true-positive associations. Using a kinship matrix as a covariate in a mixed-effect model can also help correct for population structure, but we saw very similar results using this method as using PCs.

The simplest genetic associations to disease traits involve plant pathogen resistance receptors. The gene-for-gene model of plant disease resistance was originally developed in a flax-rust system, and describes coevolution between plant resistance (*R*) genes that encode immunological resistance, and pathogen avirulence (*Avr*) genes that encode virulence factors (Flor 1956). R genes that have been successfully cloned often include nucleotide-binding leucine-rich repeat proteins (NB-LRRs; McHale et al. 2006)*. Avr* genes often encode pathogen effectors, proteins that can dismantle host defenses. Upon binding a pathogen effector protein, or a host protein targeted by an effector, the NB-LRR receptor undergoes a conformational shift to an ATP-bound state that triggers the initiation of the hypersensitive response and apoptosis of infected cells (Shao et al. 2019). While our study found outlier loci linked to several genes with LRR domains that may act as R-genes, most of the loci were linked to genes without clear associations with R-gene resistance.

An alternative strategy for resisting fungal infection is the production of antifungal specialized metabolites (Piasecka, Jedrzejczak-Rey, and Bednarek 2015). Specialized metabolites can be constitutively produced, or produced in response to receptor-binding with a pathogen-produced molecule. Production of antipathogenic compounds typically involves a greater number of genes controlling different aspects of the production, movement, and storage of specialized metabolites (Corwin and Kliebenstein 2017). More generalized pathogen resistance through metabolites could be especially selectively favored when pathogen populations evolve rapidly, since single R-genes would be quickly defeated (Hulse et al. 2023). Our results suggest a polygenic basis of adaptation, and we found several genes related to the production of specialized metabolites. Of particular note was an overrepresentation of terpenoid cyclases in genes linked to outliers in the north. We found an enrichment of copalyl diphosphate synthases, crucial intermediaries in biosynthesis pathways that produce cyclic and di-terpenoid compounds, many of which have known antifungal properties in switchgrass (X. Li et al. 2023). In maize, the gene *An2* encodes a similar copalyl diphosphate synthase that is crucial for producing antifungal phytoalexins (Vaughan et al. 2015; Ding et al. 2020). While further work will be needed to characterize these genes in switchgrass, use of metabolite-mediated resistance may be an important component of polygenic resistance in switchgrass.

One surprising result we found was that the average effect sign of GWAS associations differed for top loci across regions. In SNP calling, the most common allele in a panel is referred to as the major allele, and all other alleles as minor alleles. A trait that has a lower value in individuals with a homozygote major allele and higher value with the heterozygote minor allele would have a positive effect sign, and vice versa. Since deleterious alleles are selectively purged from populations, major alleles should be more typically associated with lower disease, resulting in a positive effect sign. GWAS in human disease follows this pattern, showing that minor alleles are more likely to contribute to higher disease risk (Kido et al. 2018). Our finding that top outliers in the north tend to have lower disease with minor alleles may indicate that alleles leading to resistance to northern rust populations have been prevented from sweeping to higher levels in switchgrass populations, possibly because they are only associated with resistance in one region, or because they are linked to other alleles that result in lower fitness.

### Outlier loci are shared with fitness and climate associations

One expectation of local adaptation is that trade-offs that lead to fitness differences across environments are produced by genes responding to multiple types of environmental stress (VanWallendael et al. 2019). When stress-response genes are linked, or when genes have pleiotropic effects, a trade-off locus can cause high fitness in a locally adapted environment, but low fitness in a foreign environment. Alternatively, conditionally neutral loci, loci that contain genes with a benefit in one environment but no fitness effect in the other, could combine to produce the pattern of local adaptation (Wadgymar et al. 2017). If trade-off loci are prevalent, we would expect that QTL for multiple environmental stressors would co-locate in the genome. We tested this assumption by uncovering overlaps between loci associated with rust resistance and loci associated with other traits such as climatic niche, biomass, and survival. We found more overlaps between outlier loci in both rust damage and other traits than would be expected if each were independently distributed throughout the genome. While this pattern does not distinguish between gene-level linkage and pleiotropy, we find support for the hypothesis that locally adaptive trade-offs can be traced to loci with responses to multiple types of stress. However, an alternate explanation is that loci associated with climate, biomass, or survival are actually rust resistance loci, since variation in rust covaries to some extent with each of these variables. Further manipulative experiments will be needed to distinguish between these possibilities.

Loci that were most strongly associated with both variation in rust severity as well as climate associations mostly contained genes involved in cellular signaling, including a MYB transcription factor and two SEC14-like genes. Since many cellular signals are shared between stress response pathways, changes in these genes can have cascading effects for numerous stress responses (VanWallendael et al. 2019). SEC14 proteins are lipid transfer proteins, involved in membrane trafficking through the Golgi body and endosome (Huang, Ghosh, and Bankaitis 2016). MYB transcription factors are numerous, but many function as crucial intermediates in responses to biotic and abiotic stress (Dubos et al. 2010). Elucidating the function of these genes in switchgrass will be an important step in understanding the genetic basis of local adaptation to biotic and abiotic stress in this species.

## Conclusion

We show that locally adapted switchgrass ecotypes are infected by a pathogen that is itself locally adapted. We predict that other systems wherein abiotic selection on the host is greater than that imposed by the pathogen may show a similar pattern. Genetic associations in different geographic regions suggest distinct physiological bases to the plant-pathogen relationship in line with local adaptation. Understanding the evolutionary drivers of host-pathogen relationships will be essential to predict disease management challenges, especially as climate change stresses plants and alters pathogen populations. For breeders focused on improving crop resistance, our results uncover potentially useful fungal disease-associated genes in switchgrass, and emphasize the challenge of relying solely on large-effect resistance genes in a fluctuating world.

## Acknowledgements

We would like to thank numerous technicians and friends that gathered data, maintained switchgrass plots, and collected rust samples for these experiments, especially T. Bortnem, P. Duberney, N. Ryan, J. Stanley, M. Carey, O. Kailing, M. Swartzendruber, S. Hofmann, T. Vugteveen, and J. Lederhouse. Gary Bergstrom, Alice MacQueen, and Kevin Myers aided in the planning of the experiment and data analysis. This material is based upon work supported in part by the Great Lakes Bioenergy Research Center, U.S. Department of Energy, Office of Science, Biological and Environmental Research Program under Award Number DE-SC0018409. We received further funding from the U.S. Department of Energy through grants DE-SC0014156 to TEJ and DE-SC0017883 to DBL. Support for this research was provided by the National Science Foundation Long-term Ecological Research Program (DEB 1832042) at the Kellogg Biological Station. A portion of these data were produced by the U.S. Department of Energy Joint Genome Institute (https://ror.org/04xm1d337; operated under Contract No. DE-AC02-05CH11231) in collaboration with the user community. The work performed by Argonne National Laboratory was supported by the U.S. Department of Energy, Office of Science, Office of Biological and Environmental Research under contract DE-AC02-06CH11357.

## Data Availability

Code used to analyze data in this manuscript can be found on github: https://github.com/avanwallendael/rust_GWAS. Rust sequencing data can be found on the NCBI SRA PRJNA1106013, and switchgrass data under PRJNA622568.

