## Supplemental Material for "Local adaptation of both plant and pathogen: an arms-race compromise in switchgrass rust"

### Supplemental figures

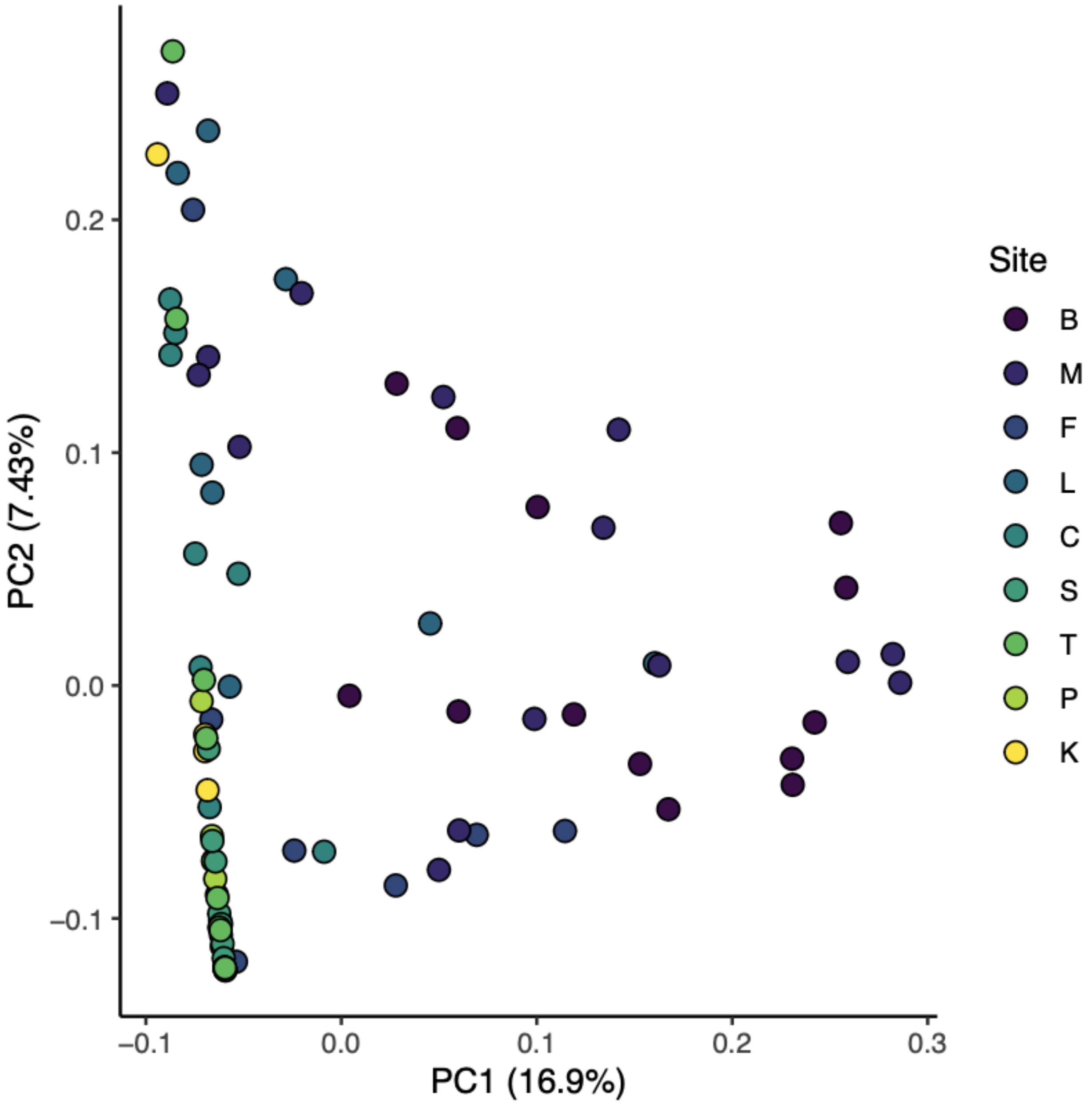


**Figure S1:** PCA of rust samples, with each point representing the rust collected from a single plant. Darker points come from higher latitudes. Axis labels indicate percent variance explained (PVE).

**A**

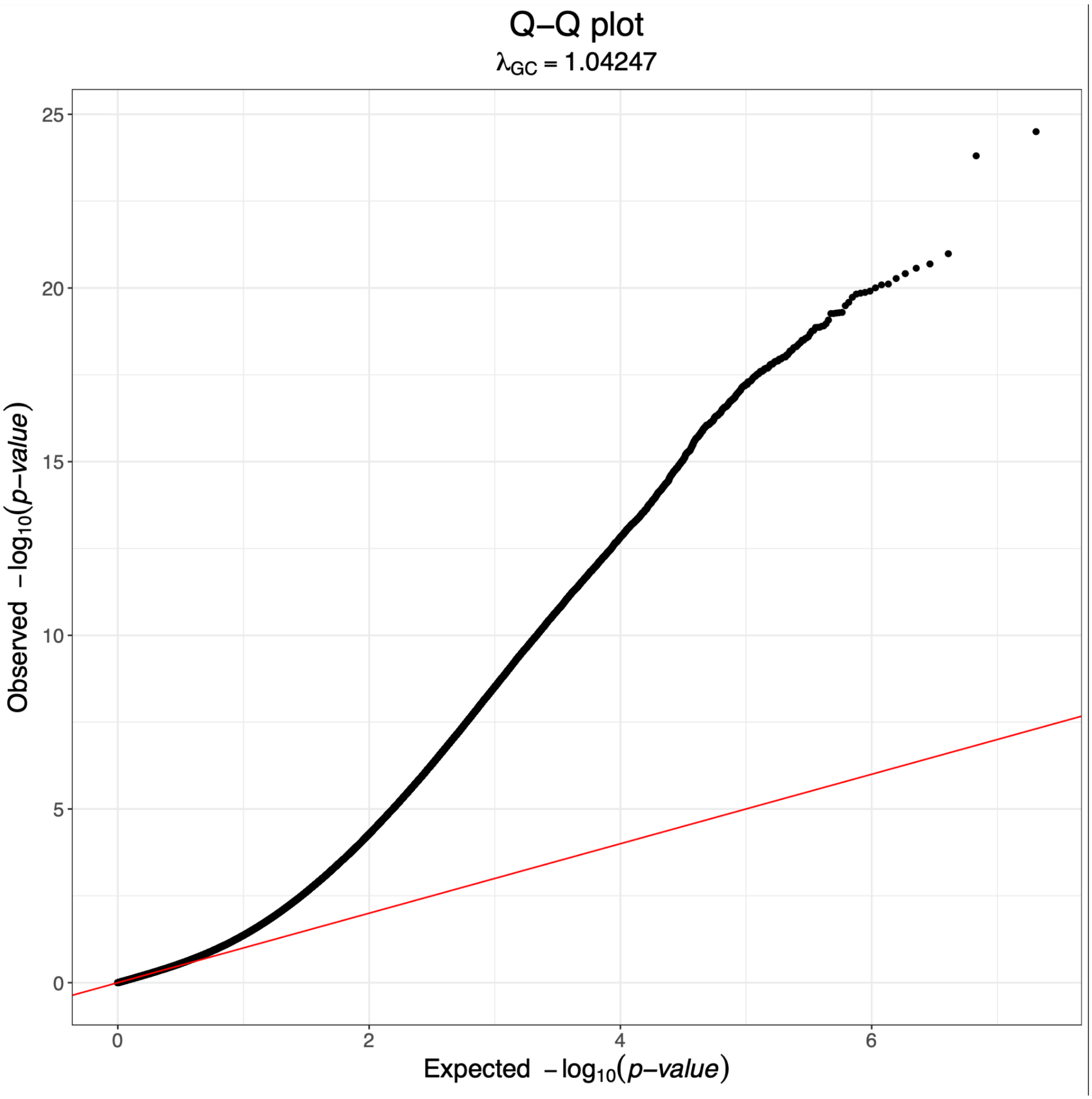

Q–Q plot
$\lambda_{GC}$ = 1.04247
25
20
15
10
5
0
0
2
4
6
Observed −log10(p−value)
Expected −log10(p−value)

B

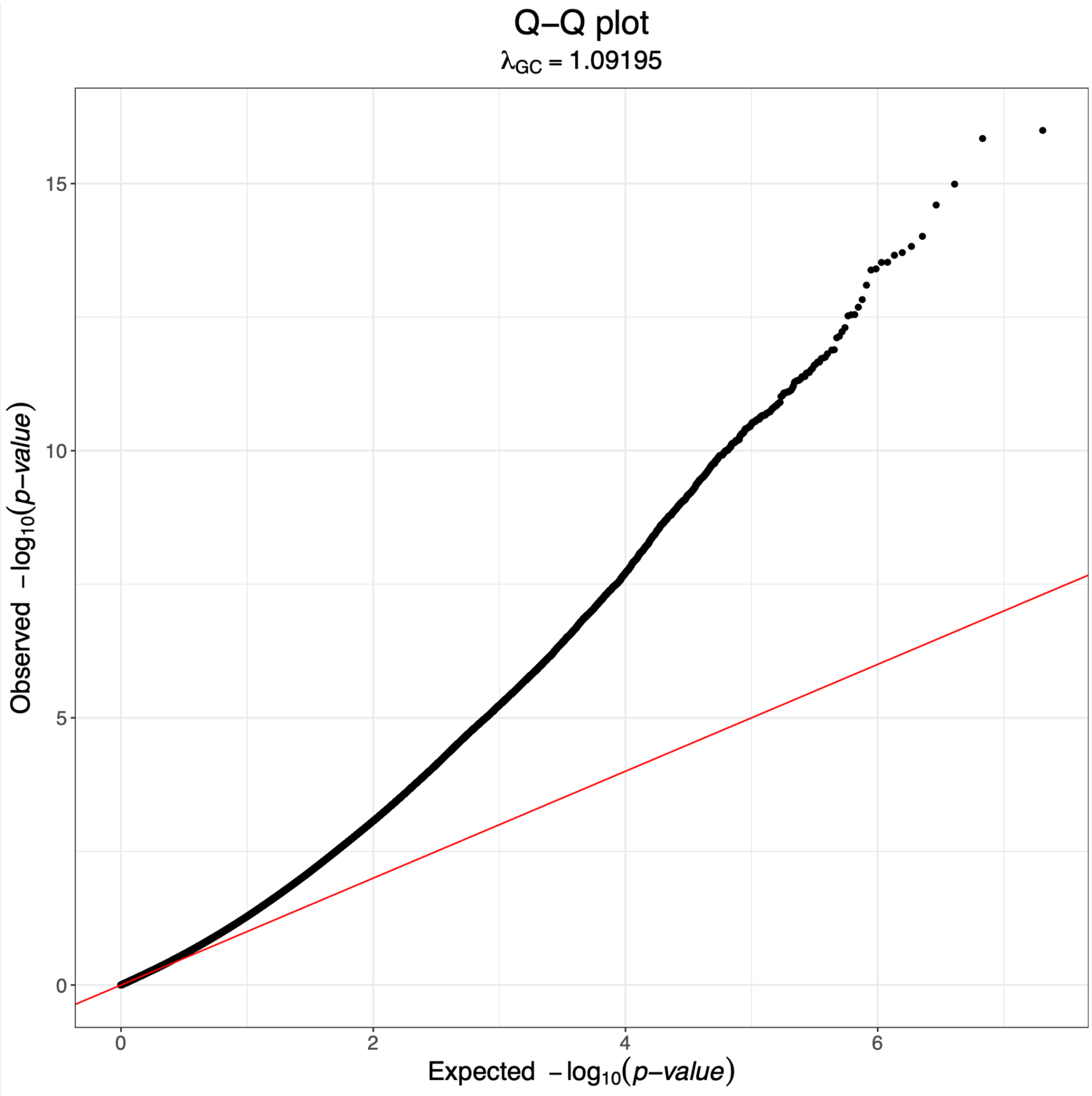


**Figure S2:** Quantile-quantile plots for northern (A) and southern (B) sites GWAS.

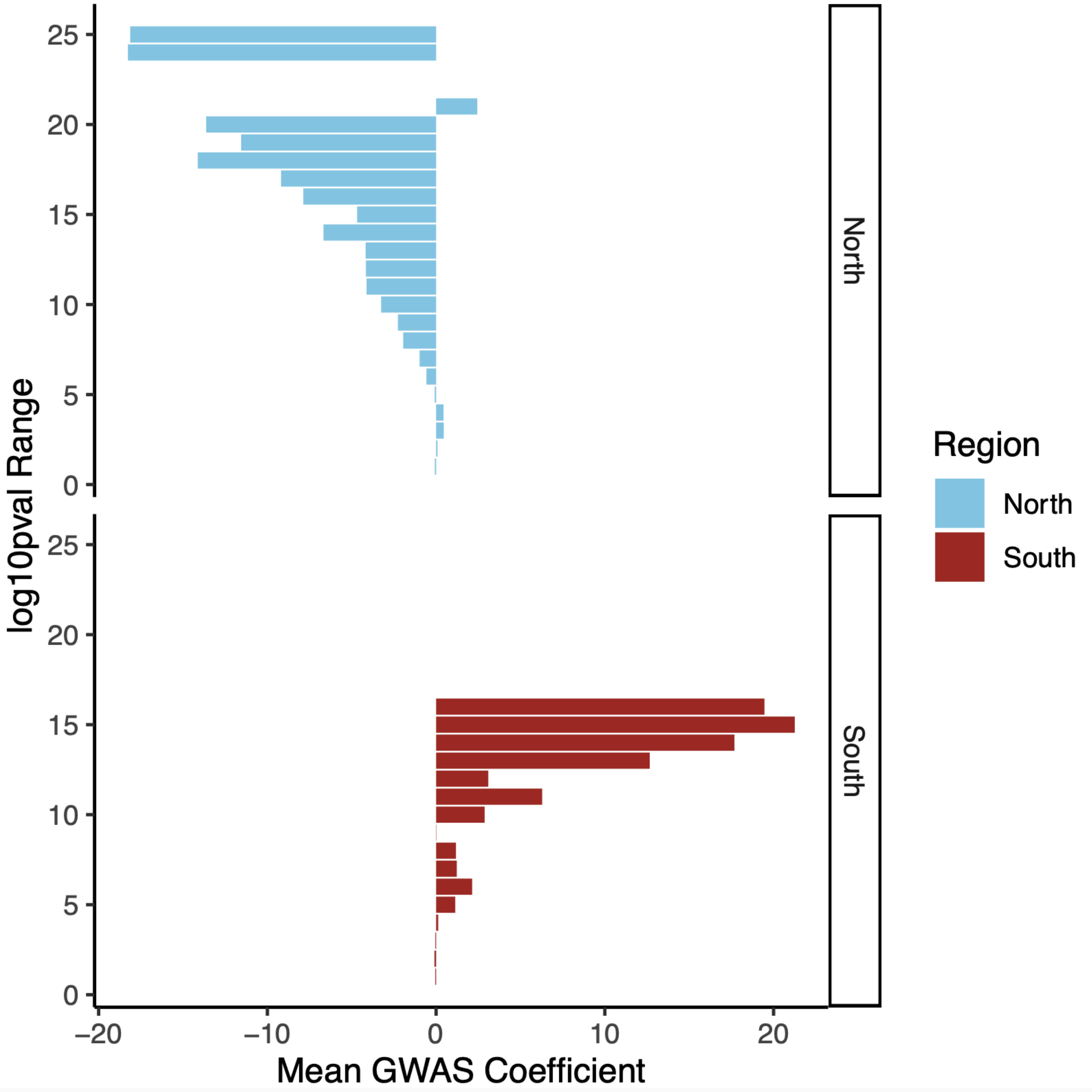


**Figure S3:** Mean coefficients for outlier loci in northern (blue) and southern (red) regions. GWAS outlier effect sizes (x-axis) were averaged for each integer range of -log10 p-value (y-axis). Higher -log10 p-values correlate positively with coefficients in the South, but negatively in the North.

**A.** North overrepresented

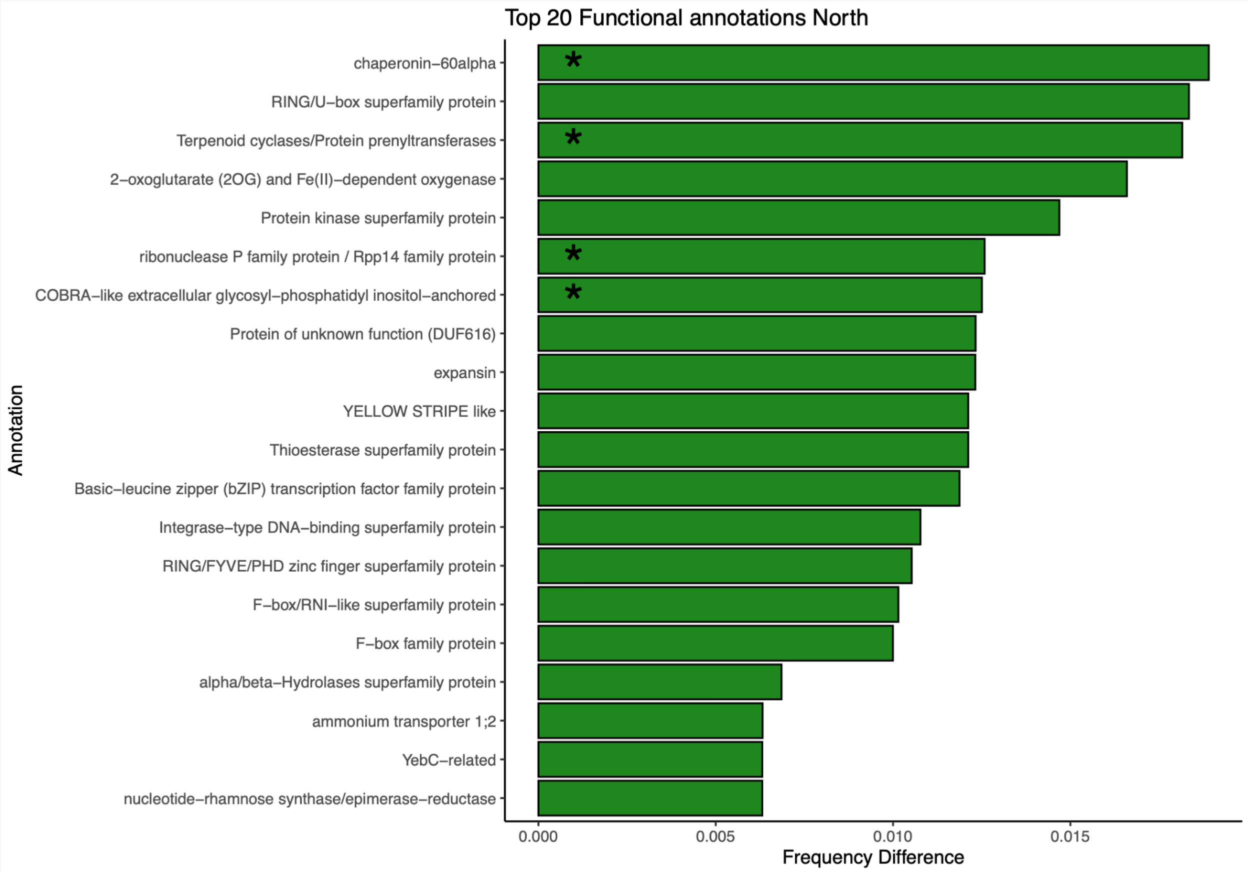


**B.** South overrepresented

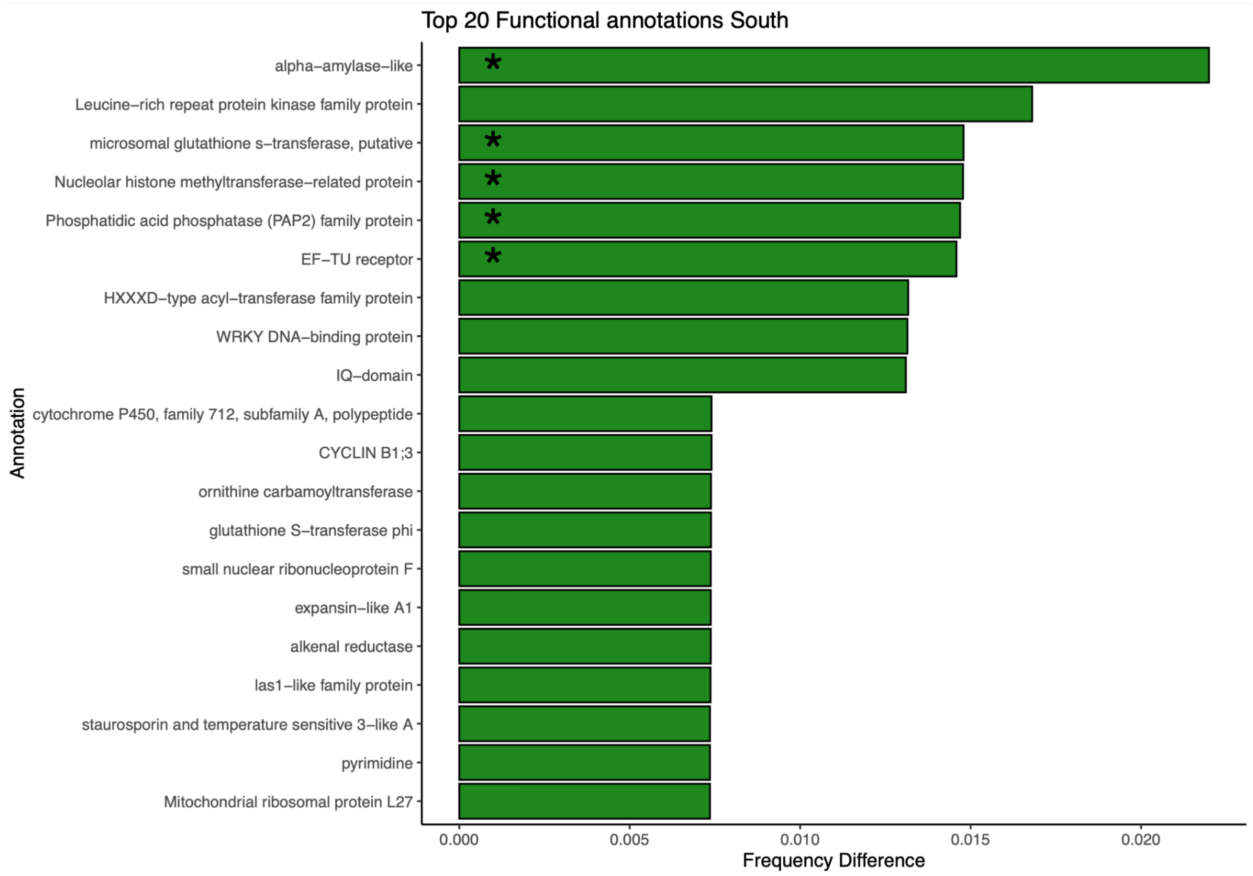


**Figure S4:** Top overrepresented gene functions linked to outlier loci in the North (**A**) and South (**B**) regions. Asterisks indicate annotations that were statistically more commonly linked to top SNPs via bootstrap resampling.

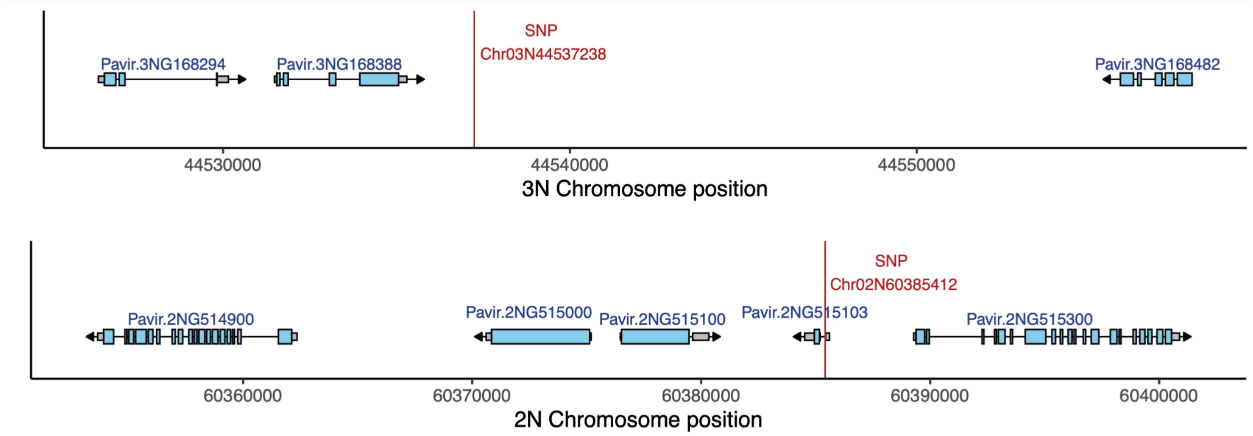


**Figure S5:** Outlier-linked regions on Chromosomes 3N and 2N. Genes Pavir.2NG514900.1, 515000.1, 515100.1, and 515300.1 have functions related to drug- or disease-resistance.

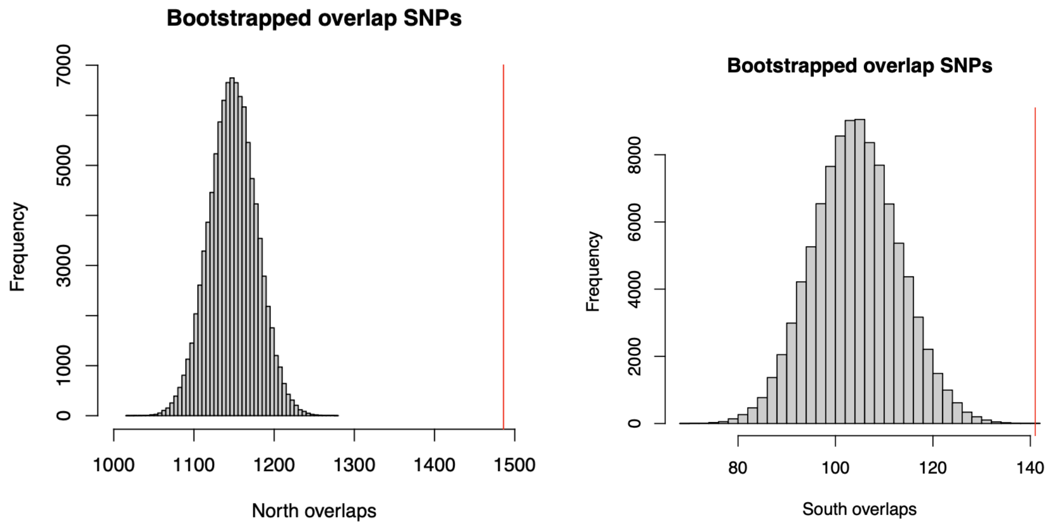


**Figure S6:** Histogram of randomly sampled loci overlapping outlier regions identified by combined trait GWAS in Lovell et al. 2021. The red line indicates the number of rust resistance SNPs overlapping combined trait outlier regions.

### Supplemental tables

Table S1: Top 20 differentially expressed genes between lowland and upland cultivars linked to GWAS outlier loci in northern and southern sites. The first column indicates the -$\log_{10}$ of an adjusted p-value from a Wald test for differential expression between ecotype.

Top DE genes North

| DE.log10p | Predicted.Function | GeneID | GWAS.log10p |
|---|---|---|---|
| 122.526074 | | Pavir.7NG105945 | 17.90188 |
| 75.143589 | RING/U-box superfamily protein | Pavir.1NG505000 | 17.45328 |
| 32.984375 | squamosa promoter-binding protein-like 12 | Pavir.5KG215300 | 20.41379 |
| 30.654452 | Terpenoid cyclases/Protein prenyltransferases superfamily protein | Pavir.1KG382200 | 20.11437 |
| 23.736580 | hydroxyproline-rich glycoprotein family protein | Pavir.2KG386500 | 18.18588 |
| 18.298091 | | Pavir.7NG018100 | 17.95064 |
| 14.275364 | Phosphoinositide phosphatase family protein | Pavir.1KG521340 | 19.26378 |
| 14.275364 | Protein kinase superfamily protein | Pavir.3NG140845 | 17.41121 |
| 14.109612 | | Pavir.2NG515103 | 23.80434 |
| 12.403643 | | Pavir.9NG187100 | 18.07645 |
| 12.313246 | ammonium transporter 1;2 | Pavir.1NG352400 | 17.22663 |
| 10.889979 | F-box family protein | Pavir.9KG451700 | 18.19136 |
| 9.302912 | peroxin 11A | Pavir.9NG683114 | 19.07573 |
| 9.092088 | Pentatricopeptide repeat (PPR) superfamily protein | Pavir.1KG506800 | 18.43072 |
| 8.883199 | | Pavir.1KG549101 | 20.56880 |
| 8.872252 | Transducin family protein / WD-40 repeat family protein | Pavir.2KG386600 | 18.18588 |
| 8.138290 | ribonuclease P family protein / Rpp14 family protein | Pavir.7NG018200 | 17.95064 |
| 8.088684 | Protein kinase superfamily protein | Pavir.9NG186800 | 18.07645 |
| 7.675674 | Terpenoid cyclases/Protein prenyltransferases superfamily protein | Pavir.1KG382115 | 17.39053 |
| 7.594359 | 2-oxoglutarate (2OG) and Fe(II)-dependent oxygenase superfamily protein | Pavir.1NG460500 | 19.91066 |

Top DE genes South

| DE.log10p | Predicted.Function | GeneID | GWAS.log10p |
|---|---|---|---|
| 24.883186 | small nuclear ribonucleoprotein F | Pavir.5NG167600 | 12.54692 |
| 20.846681 | FRIGIDA-like protein | Pavir.2KG258800 | 11.39025 |
| 18.435624 | | Pavir.3KG041145 | 10.70215 |
| 18.224244 | | Pavir.1NG427100 | 13.09818 |
| 17.369685 | Galactosyltransferase family protein | Pavir.4KG405400 | 10.89895 |
| 16.931279 | bidirectional amino acid transporter 1 | Pavir.3NG006300 | 10.65573 |
| 16.832849 | Homeodomain-like protein with RING/FYVE/PHD-type zinc finger domain | Pavir.5NG495200 | 11.10428 |
| 16.543572 | phytochrome C | Pavir.9KG462324 | 10.66392 |
| 12.987529 | | Pavir.3KG554100 | 10.90530 |
| 12.773770 | alpha-amylase-like | Pavir.3NG274900 | 12.11334 |
| 12.387746 | CYCLIN B1;3 | Pavir.5NG167700 | 12.54692 |
| 10.903161 | | Pavir.9NG458114 | 10.62876 |
| 10.693744 | RNI-like superfamily protein | Pavir.4KG405200 | 10.89895 |
| 10.639905 | alpha-amylase-like | Pavir.3NG274800 | 12.11334 |
| 10.333436 | wall associated kinase 3 | Pavir.9KG404732 | 11.46519 |
| 10.136263 | dicer-like 3 | Pavir.9NG292600 | 10.58746 |
| 10.077741 | | Pavir.6KG013176 | 10.79939 |
| 9.978394 | WRKY DNA-binding protein 40 | Pavir.2KG335400 | 13.65912 |
| 9.155404 | Malectin/receptor-like protein kinase family protein | Pavir.9KG404600 | 11.46519 |
| 9.125201 | pyrimidine 2 | Pavir.5NG543800 | 10.77314 |

Table S2: Outlier loci shared between rust resistance and fitness/climate GWAS.

**Shared outliers**

| Position | Chromosome | -log10pval | log10bf | Closest gene | Function |
|---|---|---|---|---|---|
| 50814820 | Chr01N | 14.17 | 19.04567038 | Pavir.1NG351000 | MYB-LIKE DNA-BINDING PROTEIN MYB |
| | | | | Pavir.1NG350919 | H/ACA RIBONUCLEOPROTEIN COMPLEX |
| 30622702 | Chr02K | 12.49 | 16.29773463 | Pavir.2KG198574 | E3 ubiquitin-protein ligase HUWE1 |
| | | | | Pavir.2KG198416 | none |
| 35998413 | Chr02K | 13.25 | 14.11226183 | Pavir.2KG211312 | none |
| | | | | Pavir.2KG211000 | HISTONE-LYSINE N-METHYLTRANSFERASE ATX1-RELATED |
| 25864582 | Chr02K | 12.73 | 18.97992266 | Pavir.2KG279958 | none |
| | | | | Pavir.2KG280116 | none |
| 32513574 | Chr02K | 11.02 | 10.12219096 | Pavir.2KG280600 | SEC14 RELATED PROTEIN |
| | | | | Pavir.2KG280500 | none |
| 1699386 | Chr03K | 11.01 | 10.95227979 | Pavir.3KG016900 | STRICTOSIDINE SYNTHASE-RELATED |
| | | | | Pavir.3KG016800 | 2-isopropylmalate synthase (leuA) |
| 34942439 | Chr03N | 11.85 | 24.68561921 | Pavir.3NG154800 | SEC14 RELATED PROTEIN // PATELLIN-6 |
| | | | | Pavir.3NG154900 | Glycoside hydrolase family 3 |
